# IGF1R-Targeted Delivery of a Bridged Nucleic Acid Oligonucleotide-Peptide Conjugate for MicroRNA-21 Inhibition in Triple-Negative Breast Cancer

**DOI:** 10.1101/2025.03.09.642231

**Authors:** Y.-Y. Jin, V. S. Desai, Jason Mazzaroth, E. Wickstrom

## Abstract

Triple-negative breast cancer (TNBC), defined by the absence of ER, PR, and Her2, impacts over 46,000 U.S. women annually, disproportionately affecting minority ethnic groups and individuals with BRCA1 mutations. Despite advancements such as PARP inhibitors, TNBC remains highly aggressive, with frequent recurrences and a 50% mortality rate within four years, underscoring the urgent need for more effective targeted therapies. MicroRNAs (miRNAs) represent a novel therapeutic approach. In TNBC, overexpressed miR-21 drives tumor progression, immune evasion, treatment resistance, and metastasis. Targeted miR-21 inhibition could curb these effects while minimizing harm to normal cells. We developed a peptide-conjugated miR-21 inhibitor targeting TNBC cells via the overexpressed IGF1 receptor (IGF1R), associated with poor prognosis. Using aminomethyl-bridged nucleic acid (BNA) chemistry, a serum-stable, low-toxicity anti-miR-21 RNA analog was created and tested for its effects on TNBC cell proliferation, apoptosis, tumor suppressor expression, and immune checkpoint regulation. Conjugation to an IGF1 peptide analog improved delivery, demonstrating tumor-specific biodistribution, efficacy, and safety in TNBC-bearing mice. The miR-21 inhibitor-peptide conjugate reduced proliferation, induced apoptosis, elevated tumor suppressors, and suppressed immune checkpoints in TNBC cell lines. *In vivo*, it targeted tumors, halted growth, and showed no liver or kidney toxicity, supporting its potential as a targeted, low-toxicity TNBC therapy.

## Introduction

Over 46,000 new US breast cancers every year lack 3 key drug-accessible targets: estrogen receptor, progesterone receptor, and Her2 [5]. These triple negative breast cancers (TNBC), a heterogeneous distribution of disease subtypes [7], primarily appear in younger women with African, Asian, Hispanic, or *BRCA1* mutant ancestry [5, 8, 9]. Standard of care surgery, chemotherapy, and radiation provide median survival of only 4 years [10]. While many TNBC patients may respond initially to first or second line therapy, resistance arises from mutations that drive higher oncogene expression [11, 12].

Approved molecularly-targeted drugs for TNBC include poly(ADP-ribose) polymerase (PARP) inhibitors [13], topoisomerase inhibitors [14], immunological checkpoint inhibitors [15], and antibody-drug conjugates [16], all targeting proteins. These are only modestly effective in treating small subsets of TNBC patients, such as those with *BRCA1/2* mutations [17], or elevated PD-L1 [18]. Bevacizumab and EGFR inhibitors failed to significantly improve patients’ overall survival [19]. Tecentriq, an anti-PD-L1 mAb, was initially approved upon report of one extra month of survival, but was subsequently withdrawn [20]. Trodelvy, an FDA-approved conjugate of anti-Trop-2 with irinotecan, only increased TNBC overall survival by 3 months [21]. Thus, a new molecular therapeutic agent with greater efficacy, specificity, and safety will bridge the gap in TNBC therapy. We address this critical need by targeted inhibition of oncogenic microRNAs (miRNAs) to enhance therapeutic precision and effectiveness.

miRNAs are non-coding RNAs that disrupt target mRNA translation or induce target mRNA degradation [22]. Dysregulation of miRNAs plays important roles in various stages of tumor development among different types of cancer [23–26], including breast cancer [27]. Therapeutically targeting miRNAs offers a powerful approach to overcoming cancer heterogeneity, as each miRNA can simultaneously regulate multiple mRNA networks, thereby influencing diverse cellular pathways in a coordinated manner. Most TNBC cells and adjacent stroma show high expression of microRNA-21 (miR-21) [28, 29], correlating with poor disease-free survival [28–30]. Overexpression of miR-21 was universally observed in breast cancer cell lines and tissues [31–33], boosting cell proliferation, invasion, metastasis [30, 34, 35], and drug-resistance [36, 37]. Highly metastatic 4T1 TNBC cells failed to grow tumors in miR-21 knockout Balb/c mice [38]. The mature miR-21 sequence is conserved in mice and humans.

Blocking oncogenic miR-21 can inhibit tumor progression, invasion, metastasis, angiogenesis by restoring the expression of tumor suppressor proteins, such as PDCD4 [39], PTEN [40], SPRY2 [41], maspin [42], and TPM1 [43]. PDCD4 and PTEN proteins are negative regulators of the PI3K/AKT signaling pathway. PDCD4 protein expression level was reduced in at least six human tumor types, including lung, brain, kidney, breast, colon and pancreas [44–47]. PDCD4 regulates eIF4A [48], p21, Cdk4, ornithine decarboxylase, carbonic anhydrase II and JNK/c-June/AP-1 [49]. PTEN negatively regulates cell proliferation and survival. Impairment of PTEN regulation is thought to play a role in oncogenic transformation [50]. Maspin, SPRY2, and TPM1 proteins play major roles in restricting metastasis by controlling cell branching, migration, blood vessel formation, and invasion [41, 42, 51]. Moreover, miR-21 expression can be upregulated in response to cytokine stimulation through transcriptional activation by STAT3 [52], resulting in resistance to cytokine type I interferon (IFN)-induced apoptosis [53]. This might correlate miR-21 with resistance to cell death by chemo and cytokine therapy. Since miR-21 contributes to cancer cell survival through multiple modes of action, its suppression will benefit the majority of TNBC subtypes [7]. A 19mer anti-miR-21 with particular backbone modifications has been patented for use in a liver disease [54], Alport’s syndrome, but it has not yet been applied to TNBC. Our goal is to develop a miR-21 inhibitor specifically tailored for TNBC, addressing an urgent need in combating this aggressive cancer.

To improve cancer-specific delivery of the miR-21 inhibitor, we utilize the insulin-like growth factor receptor (IGF1R) as a gateway to facilitate endocytosis. IGF1R belongs to the receptor tyrosine kinase (RTK) family, with a ligand binding domain on the cell surface, a hydrophobic transmembrane domain, and an intracellular tyrosine kinase domain followed by a C-terminal domain in the cytoplasm [55]. Upon IGF1 binding to one of the extracellular domains of the IGF1R dimer, the cytoplasmic kinase domain is activated, triggering multiple signaling pathways that drive cell division [55]. Aggressive breast cancers, including TNBC, frequently exhibit elevated IGF1R levels [56]. Approximately 50% of breast cancer [57], and 22-46% of TNBC express IGF1R [57, 58]. Most importantly, TNBC cells show strong IGF1R signaling activation, correlating with poor survival [57]. Previously, we utilized a small peptide ligand of IGF1R to selectively deliver radiolabeled peptide nucleic acid (PNA) probes for PET imaging of oncogene mRNAs in IGF1R-overexpressing cells and tumor xenografts [59]. This approach paves the way for cancer-specific delivery of therapeutic oligonucleotides to TNBC cells overexpressing IGF1R. Currently, no known signaling feedback loop has been identified between miR-21 and IGF1R [60].

Here we report the design of an RNA analog that inhibits both cell division and immune checkpoint expression by targeting miR-21 in TNBC cells. In addition, we have developed a bispecific RNA-peptide conjugate that targets IGF1R on the surface of TNBC cells, facilitating the intracellular delivery of the miR-21 inhibitor. This approach successfully blocks TNBC progression in a syngeneic mouse model.

## Results

### miR-21 inhibitor design

We incorporated the RNA analog 2’4’-BNA^NC^ (BNA) backbone modification (**Fig. 1**) in the five 5’ nucleotides and the five 3’ nucleotides to minimize hybridization-dependent and hybridization-independent toxicity, with superior hybridization affinity T_m_ >80°C. 2’4’-BNA^NC^ (BNA) is a nuclease-resistant derivative of 2’O,4’-C-methylene-bridged nucleic acid (LNA) [4, 61] (**Fig. 1**). Kinetic data revealed that the stronger binding resulted from a much slower dissociation rate constant [62]. The BNA backbone provided much higher stability in human serum [62]. *In vivo,* BNA gapmers were highly effective against proprotein convertase subtilisin/kexin type 9 (PCSK9) in low-density lipoprotein receptor (LDLR) protein regulation in Western diet-fed C57BL/6J mice [63]. BNAs displayed low liver toxicity in those mice [63].

**Fig 1.**
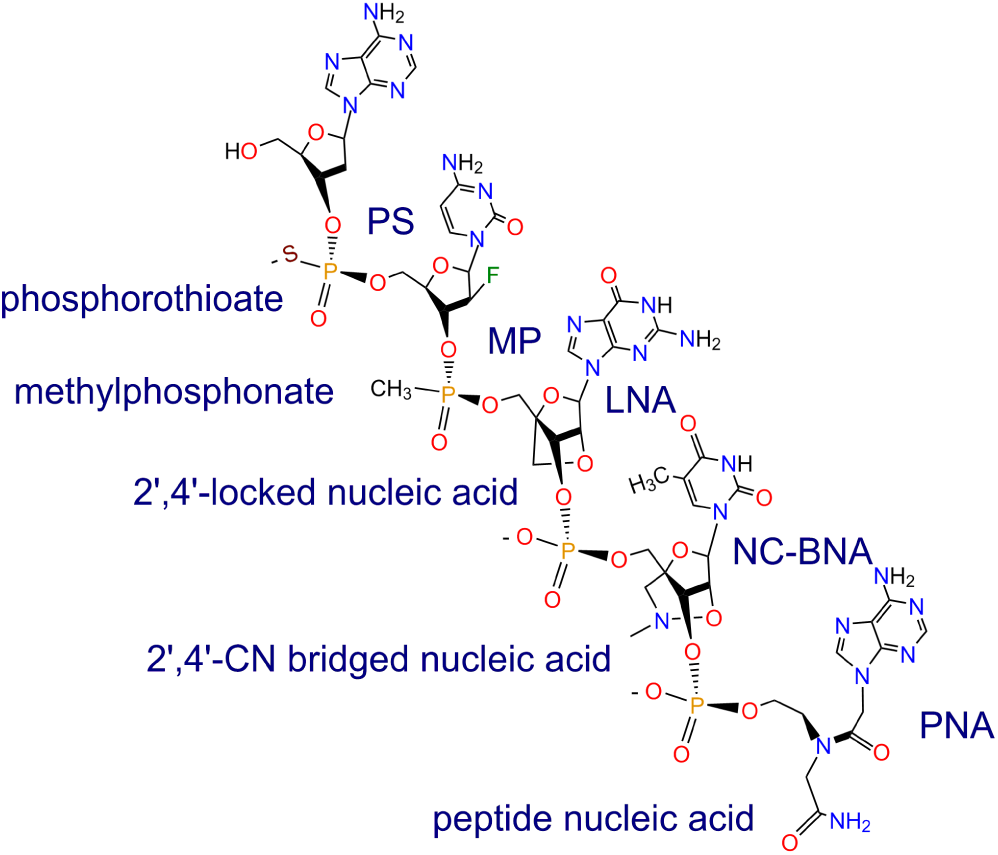
Schematics of DNA phosphorothioate (PS) [1], methylphosphonate (MP) [2], 2’-4’-locked nucleic acid (LNA) [3], 2’-NC-bridged nucleic acid (NC-BNA) [4], and polyamide nucleic acid (PNA) [6].

Whole body PET imaging of healthy rats after administration of [^18^F]BNA phosphorothioates revealed a circulation half-life ∼1 h, leaving the blood to accumulate in the liver and kidneys [64]. An anti-*PCSK9* cholesteryl-[^18^F]BNA phosphorothioate showed a shorter circulation half-life ∼20 min, with concomitant rapid accumulation in the liver, but not the kidneys [64].

BNA knockdown oligomers exhibited comparable potency to LNA [4], with minimal toxicity in mice [63]. Most recently, a BNA gapmer successfully inhibited defective CUG repeats in the 3’UTR of *DMPK* gene, which contributes to the pathogenesis of myotonic dystrophy type I (DM1) [65].

Although LNA showed greater binding affinity compared to other RNA analog modifications [66] and has been tested for therapeutic development to block miRNAs [67–69], it has been observed to induce hepatotoxicity both *in vitro* and *in vivo*, making it impractical for clinical development [70–72].

We chose a length of 15 nt to maximize uniqueness among human RNAs [73], and to minimize off-target effects [74]. All gapmer linkages are phosphorothioates. The anti-miR-21-5p sequence BND5412 (**Fig. 2**) (**Table 1**) includes a 15 nt sequence designed to specifically block miR-21-5p (identical in mice and humans). We previously reported the potential for miRNA inhibitors to mimic the effects of endogenous miRNAs [74]. We observed that conventionally designed miRNA inhibitors closely resembled the complementary passenger strand of the target miRNA within the pre-miRNA hairpin structure [74]. This finding suggests that a conventionally designed miRNA inhibitor, nearly fully complementary to the target miRNA, could mimic the function of the passenger strand in the pre-miRNA duplex, potentially leading to off-target effects. Since miRNA inhibitors often incorporate backbone modifications, the off-target effects resulting from mimicking the passenger strand could be amplified by the accumulation of these long-lasting, nuclease-resistant analogs within cells [74]. The BND5412 sequence 5’-CAGTCTGATAAGCTA-3’ was specifically selected to exclude the seed region of the passenger strand, thereby preventing the anti-miR-21 15-mer from imitating the miR-21-3p passenger strand [74].

**Fig. 2.**
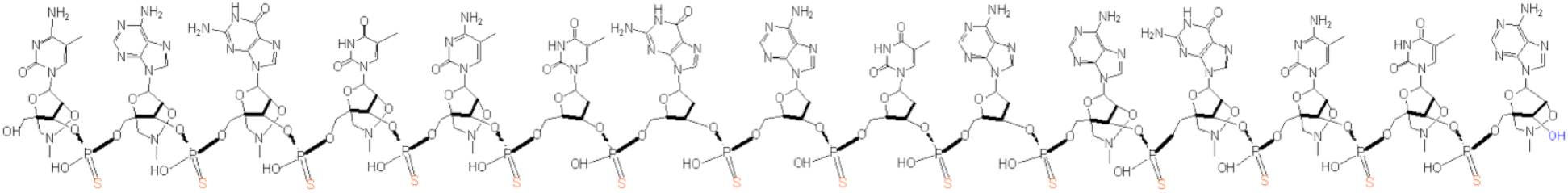
Structure of BND5412, a 15 nt miR-21 inhibitor BNA-DNA-BNA phosphorothioate. The short BNA sequence lacks the seed sequence of the corresponding miR-21 passenger strand, thus avoiding passenger strand mimicry.

**Table 1.** BNA-DNA-BNA and BNA-DNA-BNA-peptide sequences.

| Table 1. BNA-DNA-BNA and BNA-DNA-BNA-peptide sequences |  |
| --- | --- |
| Antisense Oligonucleotide | Sequence |
| Anti-miR-21 (BND5412) | 5'-**CAGTC**(TGATA)**AGCTA** |
| Scrambled (BND5371) | 5'-**TCATA**(CTATA)**TGACA** |
| Anti-miR-21-peptide conjugate (BND6482) | 5'-**CAGTC**(TGATA)**AGCTA**-click-cyclo-D(**Cys-Ser-Lys-Cys**) |
| Scrambled-peptide conjugate (BND6372) | 5'-**TCATA**(CTATA)**TGACA**-click-cyclo-D(**Cys-Ser-Ala-Cys**) |
| BNA nucleotides are shown in bold. All gapmer linkages are phosphorothioates. click = DBCO-triazolyl-K link. |  |

### The effect of miR-21 inhibitor on TNBC cells and immune checkpoints

We observed an IC50 of 22 nM for the 15-mer BNA miR-21 inhibitor BND5412 (**Fig. 2**) in suppressing miR-21 activity after transfection into MDA-MB-231 cells, leading to derepression of a luciferase reporter vector with a miR-21 complementary sequence in the 3’-UTR of the luciferase gene (**Fig. 3A**). Transfection with the miR-21 inhibitor BND5412 significantly reduced mature miR-21 levels (**Fig. 3B**) and upregulated the mRNA and protein expression of PDCD4 (**Fig. 3B, 3C**), a known direct target of miR-21 [75].

**Fig. 3.**
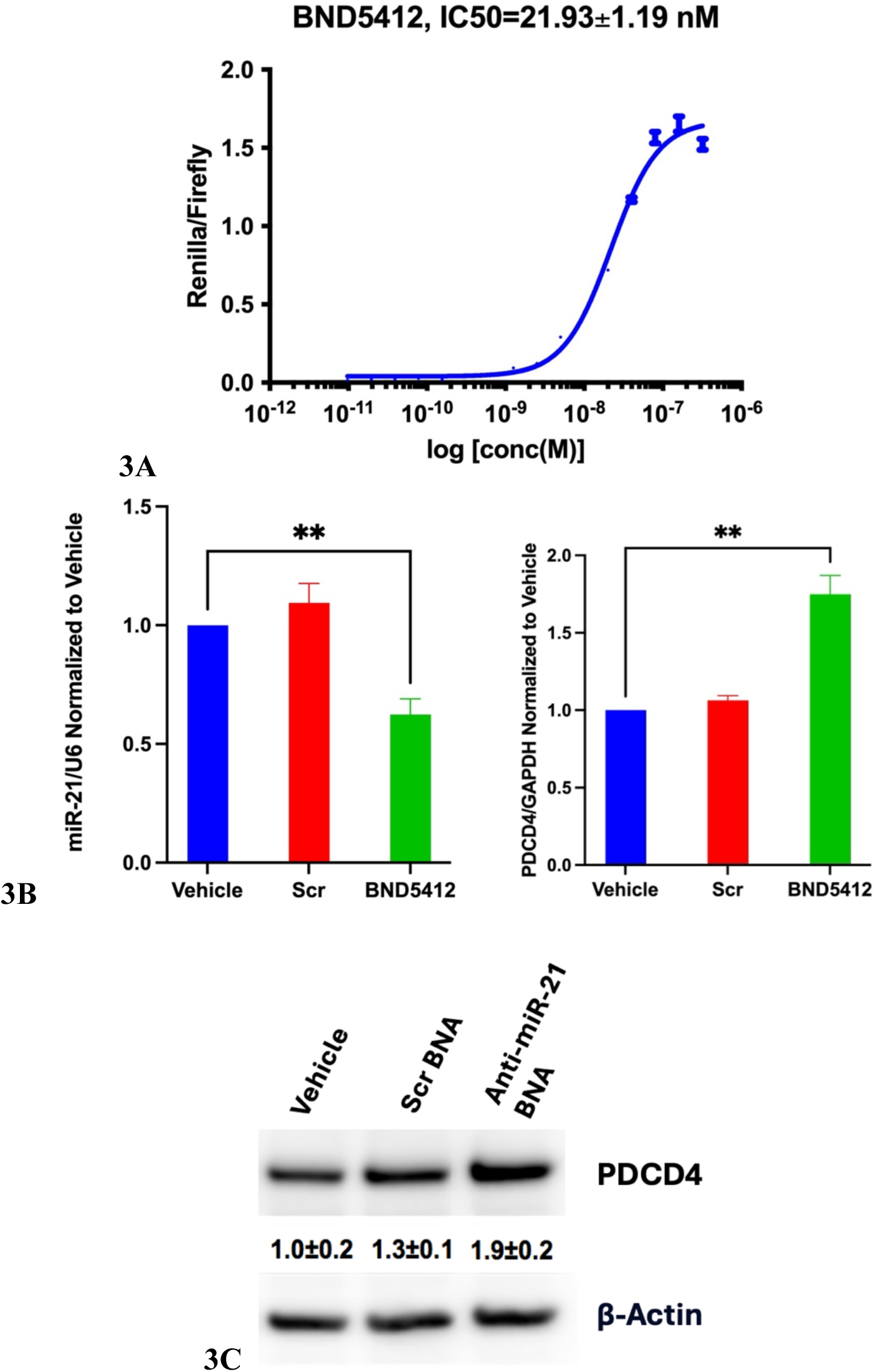
Inhibition of miR-21 activity with an anti-miR-21 BNA gamper (BND5412) in MDA-MB-231 cells. **(A)** IC50 of miR-21 inhibitor BND5412 transfected into MDA-MB-231 cells from derepression of a luciferase reporter vector with a miR-21 binding site in the 3’UTR of *Renilla* luciferase gene. Error bars, s.d. **(B)** qPCR analysis of miR-21 and *PDCD4* expression post BND5412 treatment. Error bars, s.e.m. [** = p<0.01, vs. vehicle control by 1-way t-test]. **(C)** De-repression of miR-21 direct target PDCD4 protein by western blot analysis. Triplicate mean ± s.d.

A critical aspect of miR-21’s role in the tumor microenvironment (TME) is its ability to modulate immune responses, contributing to immune evasion [76]. Suppression of miR-21 in highly metastatic tumor models led to tumor regression in immunocompetent mice, but not in immunocompromised mice. The regression was linked to increased infiltration of activated CD4+ and CD8+ T cells and a reduction in PD-L1+ monocytes, highlighting miR-21’s role in modulating immune responses [77]. Transcriptomic analysis confirmed reduced expression of cell cycle genes and enhanced immune activity upon miR-21 knockdown [77]. We investigate the effect of miR-21 inhibition by BND5412 on the suppression of several key immune checkpoint proteins. CD47, PD-L1, PD-L2, and JAK2 are critical regulators of the TME and play pivotal roles in modulating anti-tumor immunity. CD47 is a cell surface protein that binds to SIRPα on macrophages to deliver a “don’t eat me” signal, preventing phagocytosis [78]. Tumors overexpress CD47 to evade immune surveillance, hindering macrophages and dendritic cells from engulfing tumor cells [79]. PD-L1 and PD-L2 are immune checkpoints that bind to PD-1 on activated T cells, suppressing T cell function and promoting an immunosuppressive TME [80]. Tumors upregulate these ligands to evade adaptive immunity [80]. JAK2 plays a key role in cytokine signaling, essential for immune regulation and inflammation in the TME. Aberrant JAK2 activity, due to mutations or overexpression, drives production of cytokines like IL-6 and VEGF, promoting tumor growth and recruiting immunosuppressive cells such as Tregs and MDSCs [81]. This disrupts the balance between effector and suppressive immune cells, aiding tumor immune evasion [82].

Inhibition of miR-21 by BND5412 reduced the expression of immune checkpoint protein CD47, PD-L1, PD-L2, and Jak2 in human MDA-MB-231 cell line (**Fig. 4A**). Furthermore, BND5412 significantly downregulated the mRNA levels of PD-L1, PD-L2, CD47, and JAK2 in eight TNBC cell lines representing six different TNBC subtypes [83], suggesting that miR-21 blockade may promote anti-cancer immunity of TNBC cells (**Fig. 4B**). 50 nM transfection of BND5412 decreased cell proliferation by up to 8-fold in MDA-MB-231, HCC1806, BT-20, MDA-MB-157, MDA-MB-436, BT-549, and HCC1937 (**Fig. 5A**). A strong positive correlation was observed between the anti-miR-21 IC50 for inhibiting cell proliferation and miR-21 copies per cell in 6 out of 7 TNBC cell lines. The non-tumorigenic breast epithelial cell line MCF-10A was significantly less sensitive to anti-miR-21 BNA treatment (**Fig. 5B**). Furthermore, significant apoptosis, as measured by the LDH assay, was induced in TNBC cell lines MDA-MB-231, BT-20, BT-549, HCC1806, and HCC1937 (**Fig. 5C**).

**Fig. 4.**
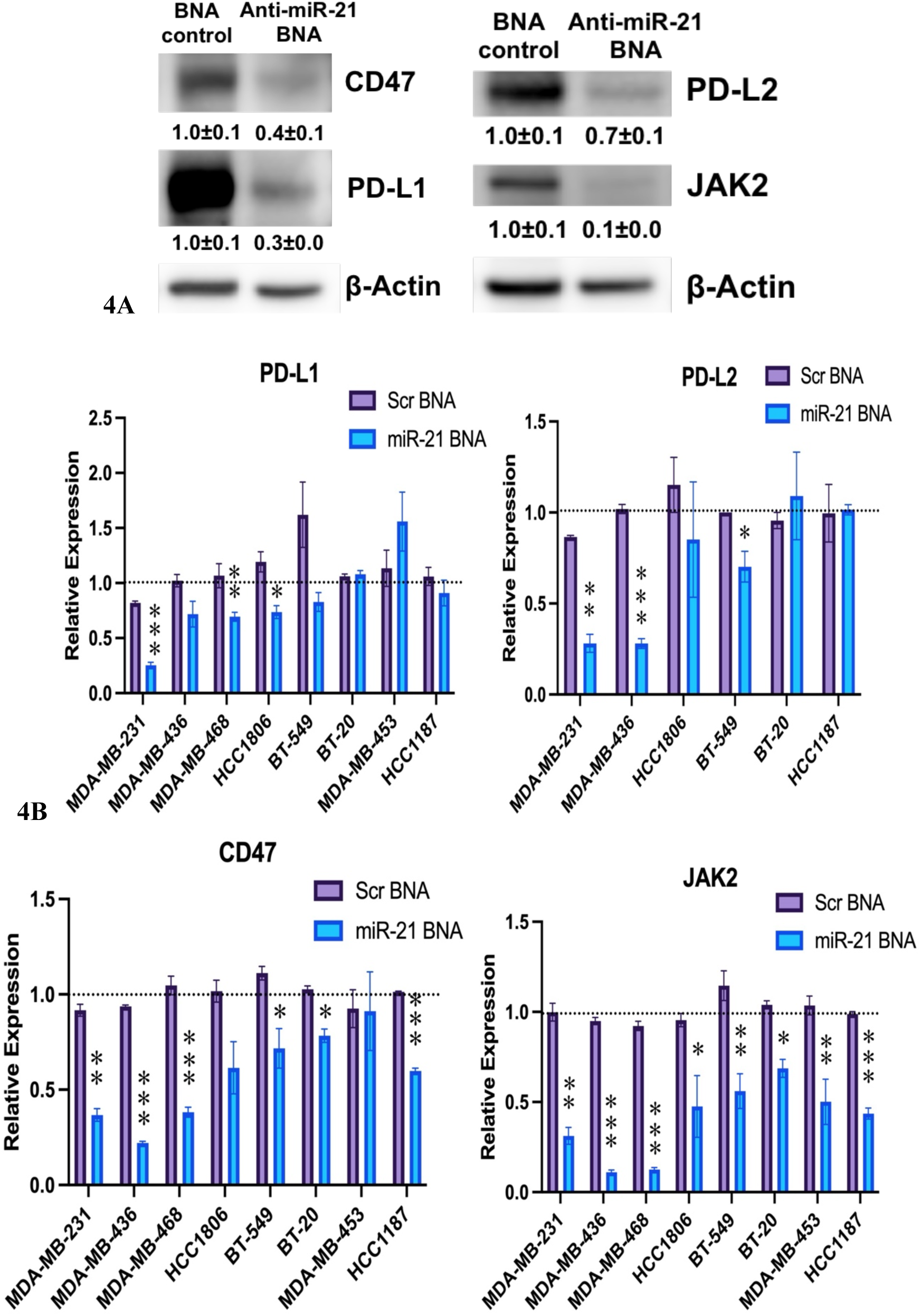
Effect of miR-21 inhibition by BND5412 on immune checkpoints in TNBC cells. **(A)** Transfection of miR-21 inhibitor BND5412 for 72 hours significantly decreased the expression of CD47, PD-L1, PD-L2, and Jak2 immune checkpoint proteins in MDA-MB-231 cells. **(B)** qPCR mRNA levels of *PD-L1, PD-L2*, *CD47,* and *JAK2* mRNA, relative to *GAPDH*, extracted from 7 TNBC cell lines 72 h post-transfection with vehicle, scrambled inhibitor (purple), or miR-21 inhibitor BND5412 (blue), in 3 biological replicates, ± s.e.m. Relative expressions were normalized to vehicle controls. [* = p<0.05, ** = p<0.01, vs. vehicle control by 1-way t-test].

**Fig. 5.**
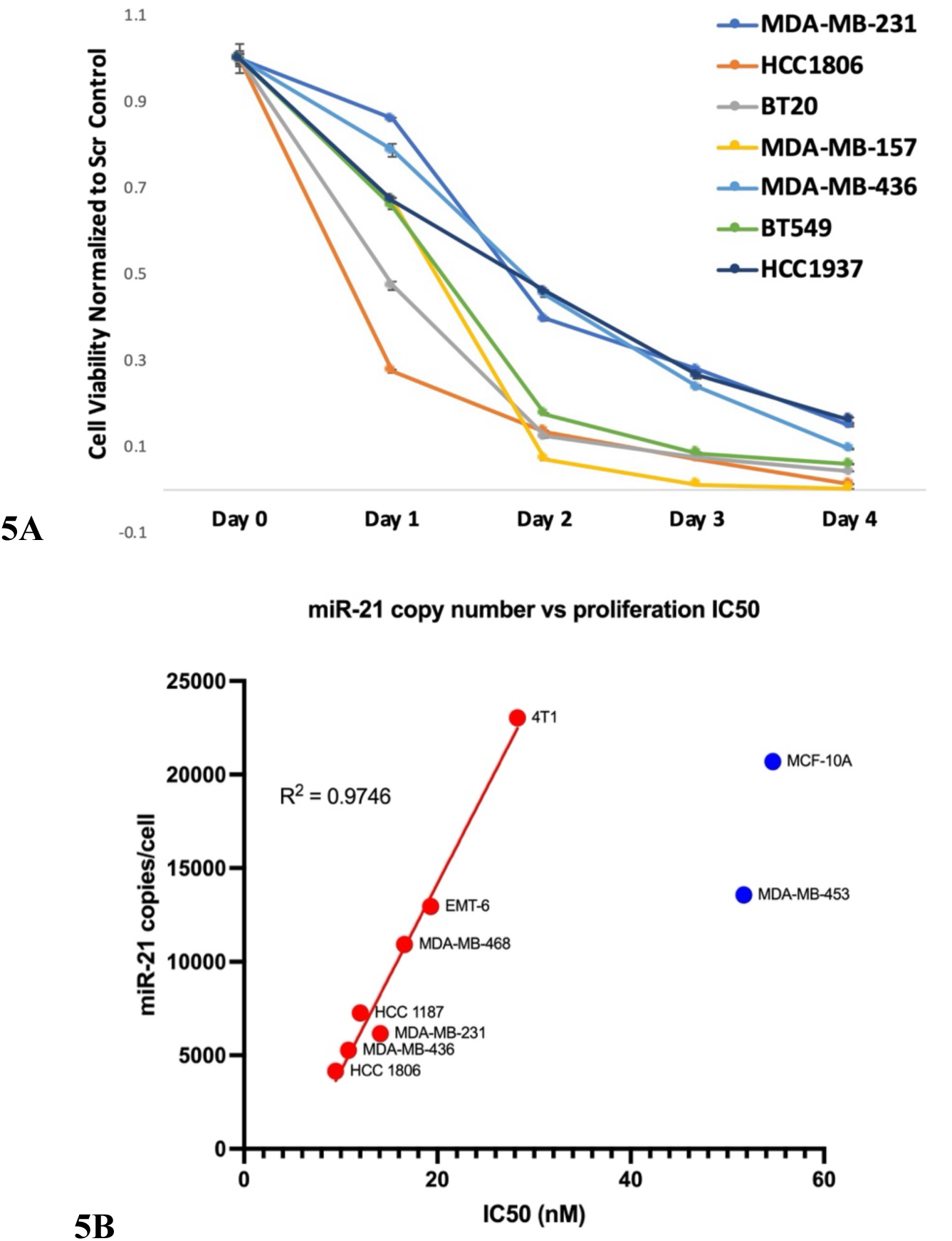
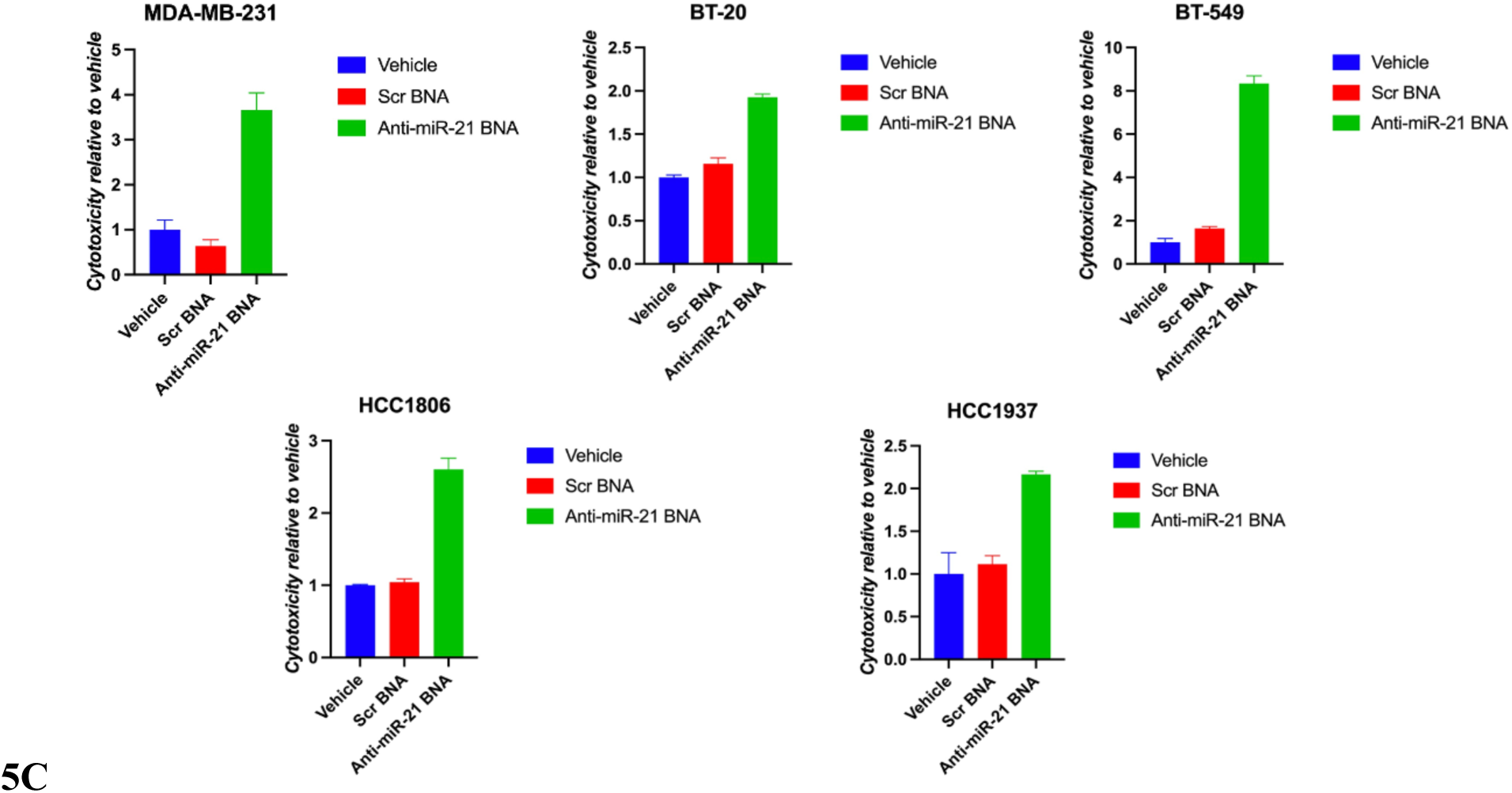
In *vitro* efficacy of miR-21 inhibitor BND5412 in TNBC cells. **(A)** Cell Titer Glo assay ± s.d. following 50 nM miR-21 inhibitor BND5412 transfected into 7 different TNBC cell lines on day 0. **(B)** Correlation between cell proliferation mean IC50 over 48 h and miR-21 mean copies/cell in 8 TNBC cell lines transfected with concentration gradients of miR-21 inhibitor BND5412. **(C)** LDH assay ± s.d. 72 h post 50 nM miR-21 inhibitor BND5412 transfection to test for apoptosis.

### RNA expression profile post miR-21 blockade

We conducted RNA-seq analysis on RNA samples from HCC1806 cells transfected with the EC90 concentration of 10.7 nM, determined from the luciferase reporter assay of the miR-21 inhibitor BND5412, alongside vehicle-treated cells. Off-target effects were assessed using GGGenome (https://gggenome.dbcls.jp) by analyzing human spliced RNAs complementary to BND5412 sequence with 0-2 mismatches, insertions, or deletions. Over 300 genes were identified in the search results.

Notably, no transcripts with 0-1 mismatches were downregulated by at least 2-fold. However, 8 genes with 2 mismatches to the miR-21 BNA sequence showed a decrease of at least 2-fold **(Fig. 6A)**. Among these, SH3PXD2A, DIAPH2, PTPRK, MGAT5, and NLGN4X were identified as oncogenes. The remaining genes, ERC1, ATRN, and FHOD3, were not linked to any known disease when downregulated. No gene exhibited off-target effects greater than a 4-fold change.

**Fig. 6.**
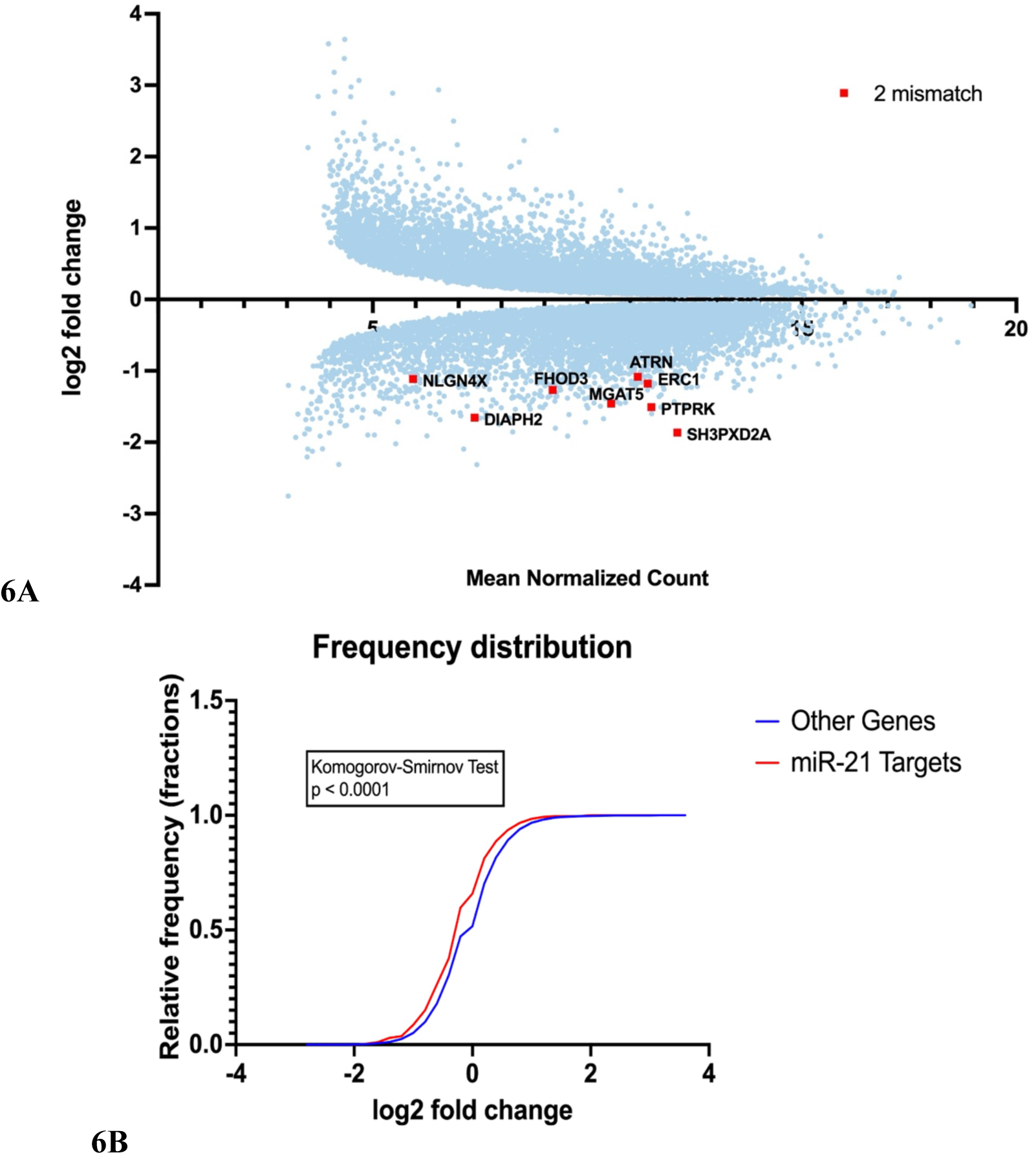
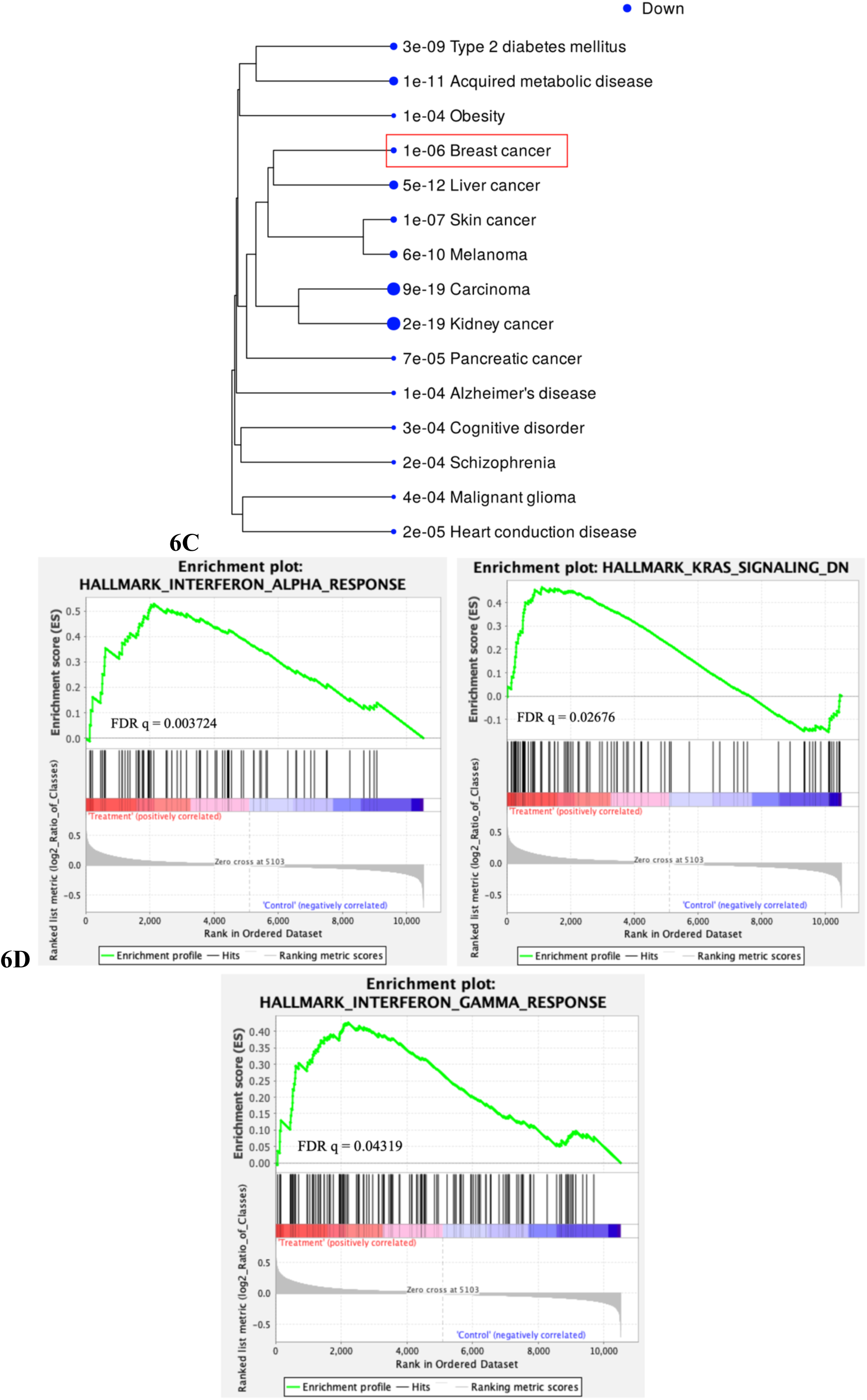
Transcriptome analysis of miR-21 inhibitor BND5412 in TNBC cells. **(A)** MA plot of genes with significant differential expression (blue), in RNA-seq analysis from 6 biological replicates of HCC1806 TNBC cells treated for 24 h with EC90 of miR-21 inhibitor BND5412. Genes exhibiting a minimum of a 2-fold change and containing 2 mismatches relative to BND5412 are highlighted in red. **(B)** Cumulative frequency distribution of miR-21 target genes (red) and all other genes with significant differential expression (blue), in RNA-seq analysis from 6 biological replicates of HCC1806 TNBC cells treated for 24 h with EC90 of miR-21 inhibitor BND5412 [Komogorov-Smirnov test, p<0.0001]. **(C)** Top downregulated disease pathways identified through Jensen Disease enrichment in HCC1806 cells treated for 24 hours with the EC90 miR-21 inhibitor in luciferase reporter assay. **(D)** GSEA analysis of top 3 enriched pathways of differentially expression genes from HCC1806 TNBC cells treated for 72 h with EC90 of miR-21 inhibitor BND5412.

To assess the global impact of miR-21 on regulated transcripts, a list of predicted miR-21 target genes was obtained from TargetScan (https://www.targetscan.org/vert_80/). A Kolmogorov-Smirnov test comparing the cumulative distribution of transcripts between miR-21 targets and non-target genes revealed a significant difference **(Fig. 6B)**. This suggests that BND5412 treatment exerted widespread on-target effects on miR-21-regulated transcripts.

Gene Set Enrichment Analysis on HallMark genes from MSigDB dataset (GSEA, https://www.gsea-msigdb.org/gsea/index.jsp) on differentially expressed genes identified top 3 enriched pathways (**Fig. 6D**). The enrichment results showed that miR-21 inhibition by BND5412 activated interferon α and interferon γ response, both of which have a positive effect on anti-tumor immunity [84, 85]. In addition, the *KRAS* inhibitory pathway was also significantly enriched in BND5412 treated samples (**Fig. 6D**). Disease enrichment analysis using the Jensen Disease database on the iDEP platform (http://bioinformatics.sdstate.edu/idep96/) revealed significant enrichment of breast cancer among down-regulated genes (**Fig. 6C**). Other top-enriched cancers included kidney, liver, melanoma, pancreatic cancer, and malignant glioma. Enrichment of diseases such as Alzheimer’s disease, schizophrenia, heart conduction disease, obesity, and metabolic disorders like type 2 diabetes was also observed. No disease association was enriched among upregulated genes. These results indicate that miR-21 inhibition may have therapeutic potential across a range of diseases beyond cancer.

### miR-21 inhibitor-peptide conjugate for cell-specific delivery

To deliver therapeutic miR-21 inhibitor to TNBC cells that overexpress IGF1R, we conjugated a small peptide analog of IGF1 [86] to BND5412 via a non-cleavable DBCO linker to endocytose the miR-21 inhibitor preferentially into TNBC cells (**Fig. 7B**) [87, 88]. To evaluate the serum stability of BND6482, 2 µM of the compound was incubated in 50% and 90% fetal bovine serum (FBS). At various time points, aliquots were collected and analyzed on a denaturing 15% TBE-Urea gel. The results showed that the majority of BND6482 remained intact after 48 hours at 37°C in both serum conditions, indicating high stability. Furthermore, the detached oligonucleotide exhibited no signs of degradation or smaller fragments, suggesting high serum stability of the 15-mer BNA gapmer (**Fig. 7C**). 100 nM Cy5-labeled miR-21 inhibitor-peptide conjugate (BND6482) demonstrated efficient cytoplasmic uptake in HCC1806 cells, as observed through live-cell confocal microscopy after 4 hours of incubation **(Fig. 8A)**.

**Fig. 7.**
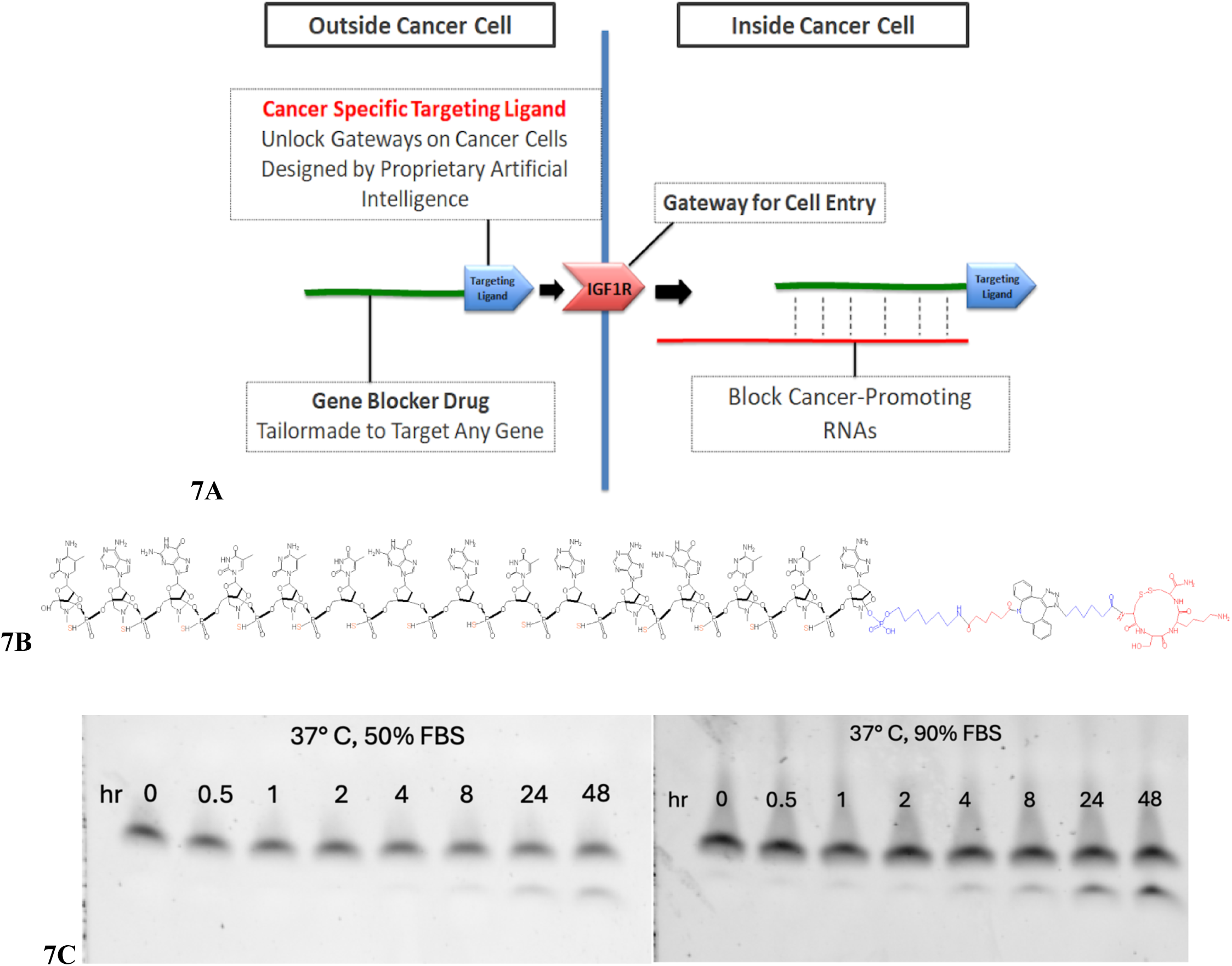
(A) Design of miR-21 inhibitor conjugate of BNA-DNA-BNA gapmer with IGF1 peptide for endocytosis by cell surface IGF1R, highly expressed on TNBC cells. **(B)** Structure of BND6482, the miR-21 inhibitor BNA-DNA-BNA phosphorothioate coupled to an IGF1 peptide analog for IGF1R-mediated endocytosis. **(C)** 15% TBE-Urea gel showing serum stability of BND6482 at 37**°**C in 50% or 90% FBS.

**Fig. 8.**
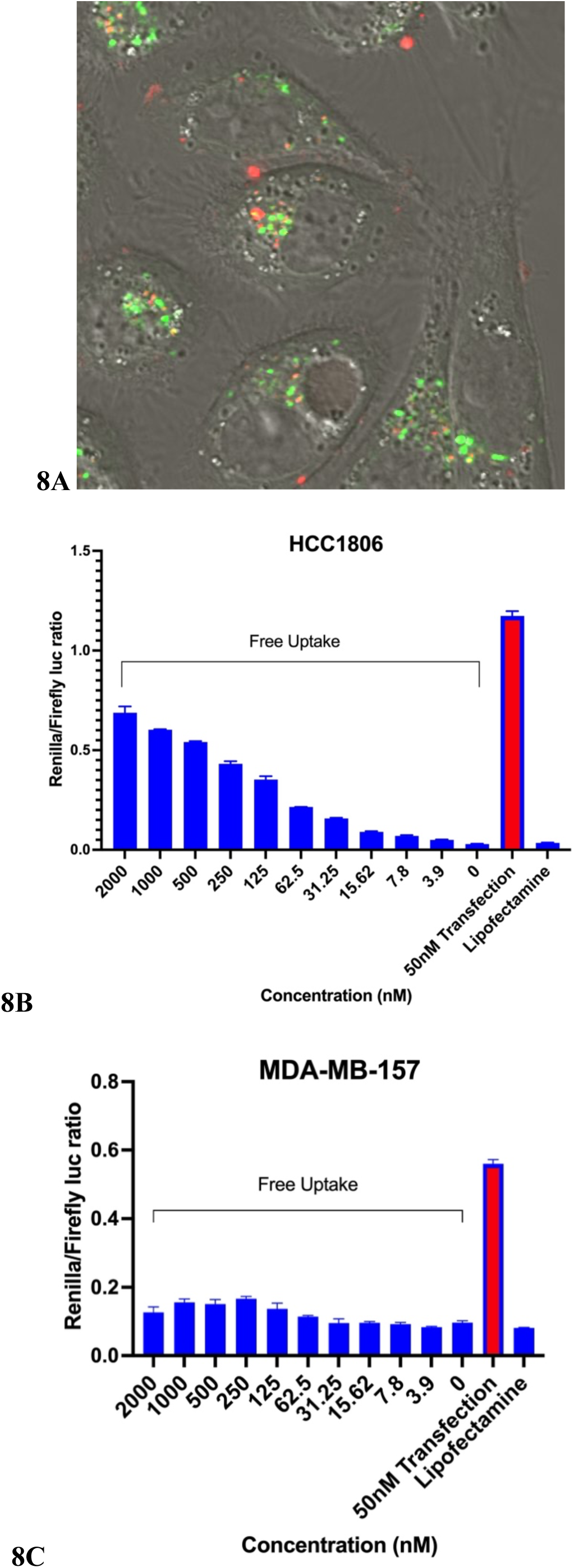
Uptake, *in vitro* specificity, and target engagement of BND6482 in TNBC Cells. **(A)** Live-cell confocal fluorescence images of HCC1806 cells after 4 h in 100 nM Cy5-BND6482. Green: LysoTracker. Red: Cy5. Yellow: Cy5-LysoTracker overlap. **(B-C)** Dose-dependent inhibition of miR-21 by BND6482 after 24h without lipofection (blue) was measured in miR-21 luciferase reporter assay in comparison to BND6482 transfected at 50 nM (red) in high IGF1R expressing HCC1806 cells **(B)**, and low IGF1R expressing MDA-MB-157 cells **(C)**. Error bars, s.d.

Fluorescent BND6482 partially co-localized with green acidic vesicles, indicated by yellow puncta, while a portion was observed outside these vesicles, suggesting partial endosomal escape. BND6482 uptake in high IGF1R-expressing HCC1806 cells [89] resulted in dose-dependent derepression of miR-21 luciferase reporter expression **(Fig. 8B)**. In contrast, uptake in low IGF1R-expressing MDA-MB-157 cells [89] did not significantly derepress the luciferase reporter without lipofection **(Fig. 8C)**.

### BND6482 biodistribution and uptake in EMT6 allografts

A single intraperitoneal injection of AlexaFluor647-BND6482 at 5 mg/kg in EMT6 allografts in immunocompetent female Balb/c mice demonstrated fluorescent drug distribution from the injection site (left) to the tumor site (right), becoming visible within 2 hours and persisting for at least 96 hours. Notably, no significant accumulation was observed in the liver or kidneys during this time **(Fig. 9A**.

**Fig. 9.**
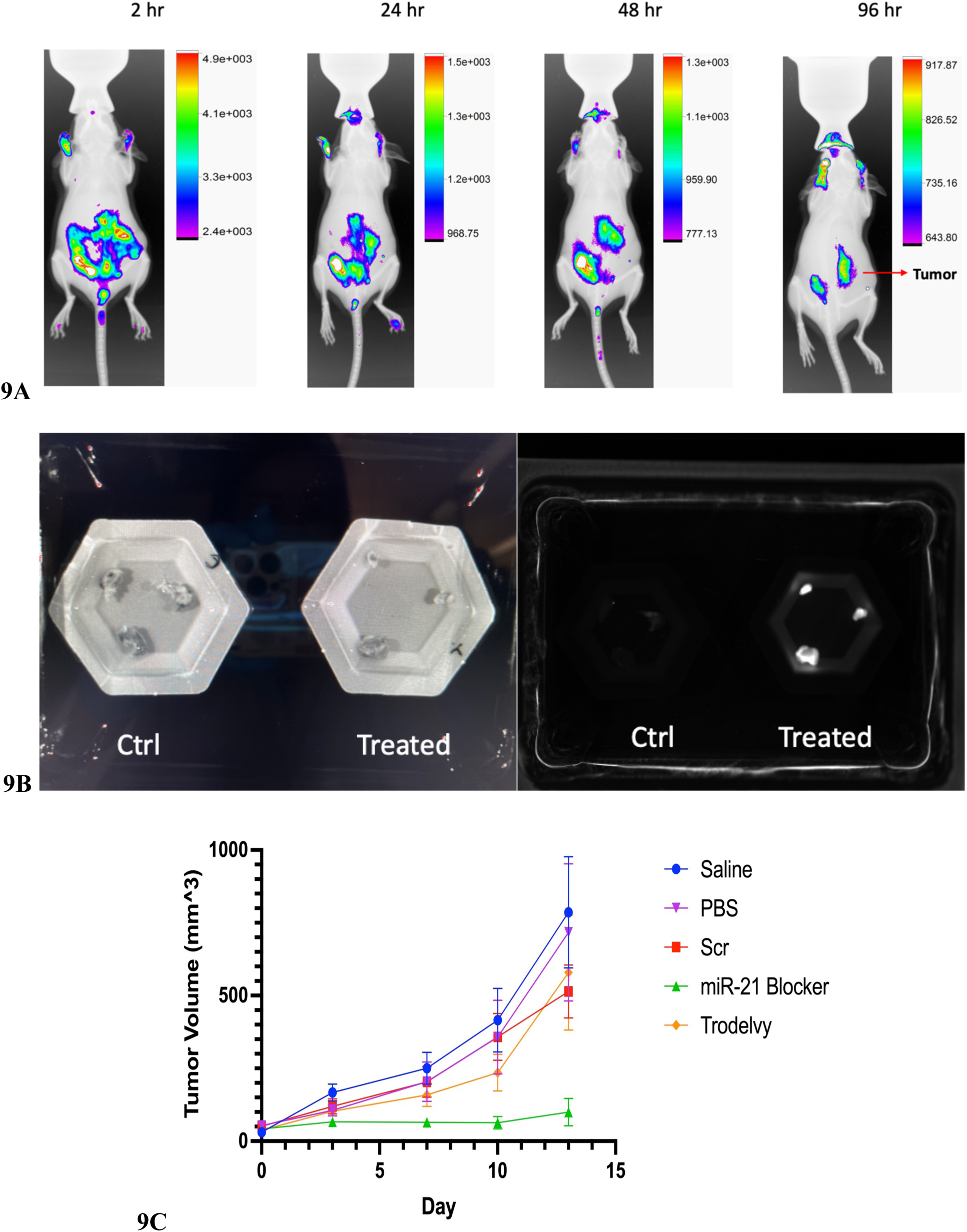
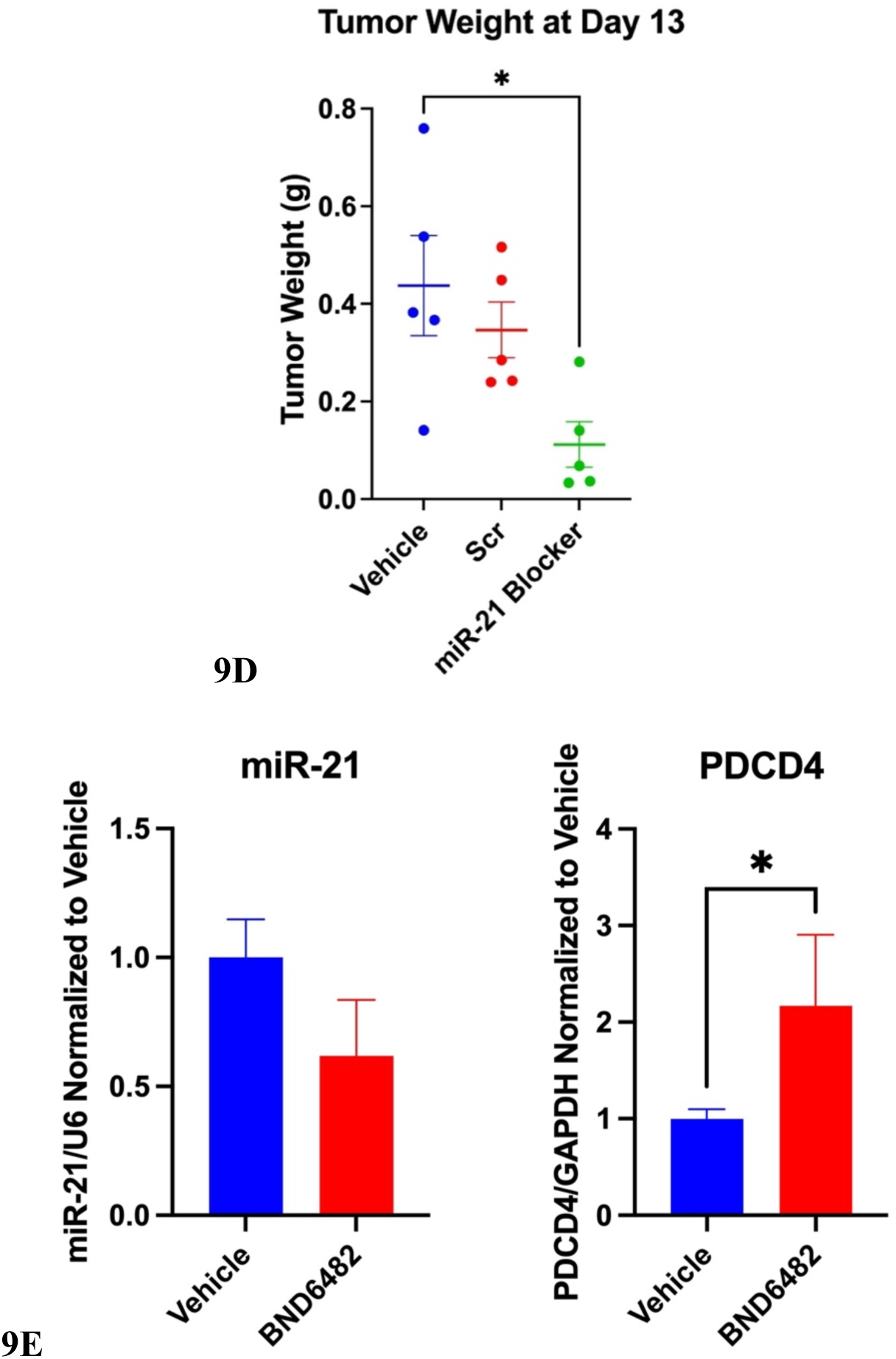
Biodistribution and therapeutic effect of BND6482 in orthotopic EMT6 allografts. **(A)** Fluorescent BND6482 distribution to EMT6 allografts in female immunocompetent female Balb/c mice 2 hr - 96 hr after a single IP injection of miR-21 inhibitor at 5 mg/kg. **(B)** Syngeneic EMT6 cells are used to generate orthotopic allografts in female immunocompetent female Balb/c mice. IP injection of saline vehicle control or AlexaFluor647-labeled BND6482 at 5 mg/kg once daily for 3 days inhibited tumor growth, as seen in white light images (left) or fluorescent images (right) of dissected tumors. **(C)** Tumor volumes of EMT6 orthotopic allografts stayed small after twice a week IP injection of 5 mg/kg of BND6482. Vehicle, scrambled drug and Trodelvy allowed continued growth. Error bars show mean with s.e.m. **(D)** Tumor masses of EMT6 orthotopic allografts were significantly reduced after twice a week IP injection of 5 mg/kg of BND6482. Error bars show mean with SEM. *p<0.05 by One-way ANOVA with Dunnett’s multiple comparison test. **(E)** qPCR measurement of miR-21 and PDCD4 mRNA after twice a week IP injection of 5 mg/kg of BND6482. n = 5; error bars, s.e.m. *p<0.05 by Mann-Whitney one-tailed t-test.

### TNBC allograft response to BND6482

Tumor inhibition was observed following daily intraperitoneal dosing of 5 mg/kg AlexaFluor647-labeled BND6482 for 3 days **(Fig. 9B)**, with two out of three tumors markedly smaller than those in the vehicle group. Fluorescent imaging confirmed drug accumulation in the extracted tumors from each treated animal **(Fig. 9B)**. Subsequently, we conducted a 2-week trial with nonfluorescent BND6482. After EMT6 tumor allografts reached approximately 5 mm in diameter, mice were treated with saline or PBS vehicle, 5 mg/kg Trodelvy, scrambled BND6372, or BND6482, administered intraperitoneally twice a week on days 0, 3, 7, and 10 over 13 days. Trodelvy, an FDA-approved anti-Trop-2 and irinotecan conjugate, was included as a comparison at the same dosage. Tumor volumes were measured at each injection point, and on day 13, tumors were harvested and weighed for analysis. BND6482-treated tumors stayed small without apparent tumor progression over 2 weeks (**Fig. 9C**). After 2 weeks of treatment, BND6482-treated tumors exhibited a significant reduction in tumor mass, with a 74% mean reduction compared to vehicle, Trodelvy, or scramble-treated groups **(Fig. 9D)**. Furthermore, BND6482 treatment resulted in a decrease in miR-21 level and a corresponding upregulation of PDCD4 mRNA, a known miR-21 target **(Fig. 9E)**.

### Liver and kidney toxicity assessment of BND6482

To assess liver and kidney toxicity of BND6482 at the therapeutic dose, immunocompetent female Balb/c mice bearing EMT6 tumors were treated with three intraperitoneal injections of 5 mg/kg BND6482 over a 2-week period on days 0, 6, and 12, once the tumors reached 5 mm in diameter. Blood serum was collected on day 14 and analyzed for hepatotoxicity markers, including liver enzymes (ALP, ALT, AST), bile acids, blood urea nitrogen (BUN), creatinine, and γ-glutamyl transferase (GGT). Compared to the vehicle and scrambled control groups, BND6482-treated mice showed no significant alterations in serum markers indicative of hepatotoxicity **(Fig. 10A)**. Liver and kidney toxicity in treated mice was further evaluated using fluorescent immunohistochemistry to analyze markers including caspase 3, Ki67, and IBA1. Elevated caspase 3 levels indicate drug-induced cell death in the liver and kidney, signaling toxicity [90]. Increased Ki67 expression suggests regenerative proliferation of liver cells in response to damage [90]. IBA1 serves as a marker for activated macrophages, with higher levels reflecting an early immune response to toxic substances, tissue damage, and inflammation [90]. LNA-containing antisense oligonucleotides with high toxicity was previously shown to significantly induce caspase 3, Ki67, or IBA1 levels in liver and kidney tissues of rats [90]. Liver and kidney sections from mice treated with BND6482 exhibited no significant increase in caspase 3, Ki67, or IBA1 levels compared to saline-treated controls. This indicates that BND6482 does not induce observable toxicity at the effective dose **(Fig. 10B)**.

**Fig. 10.**
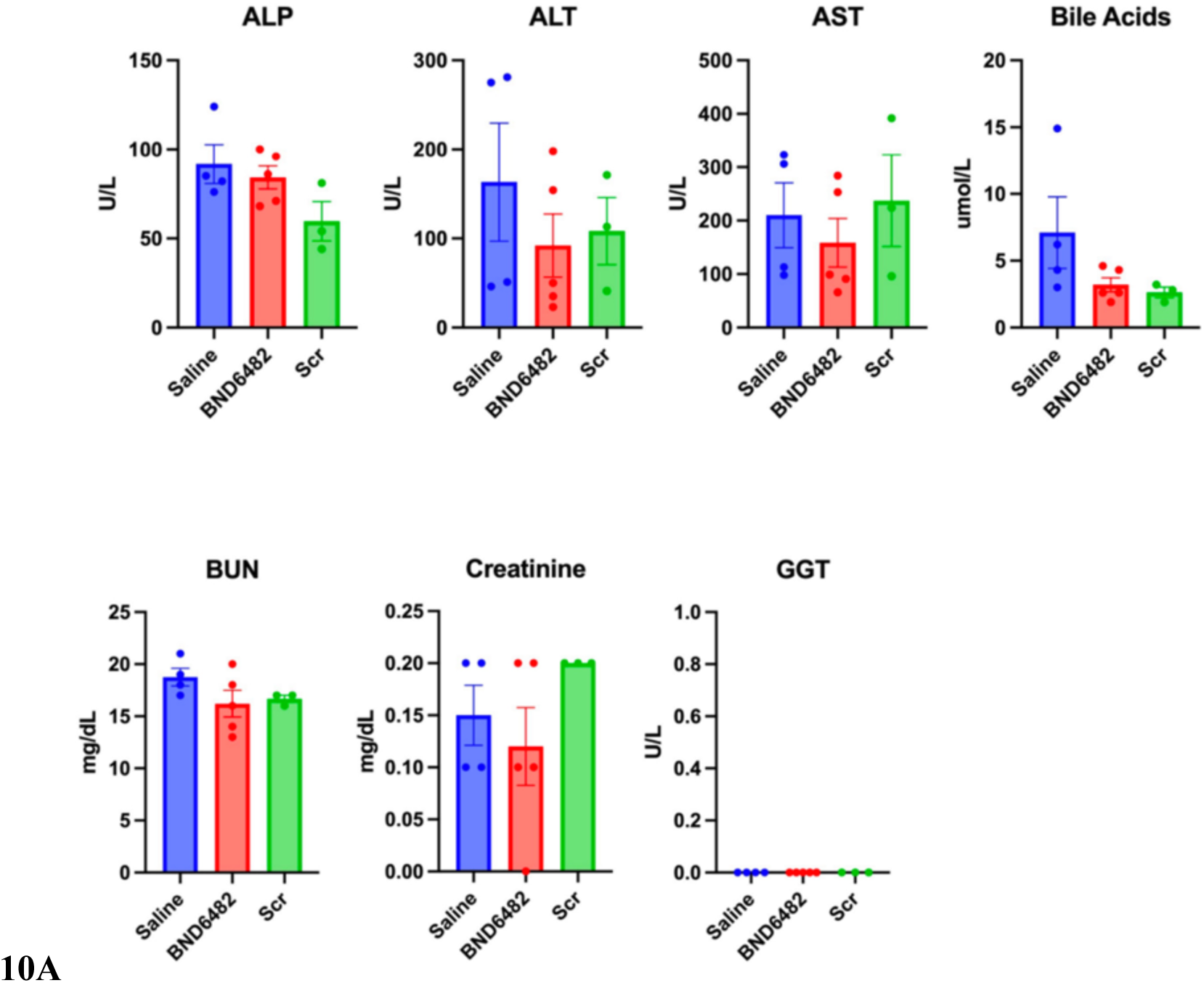
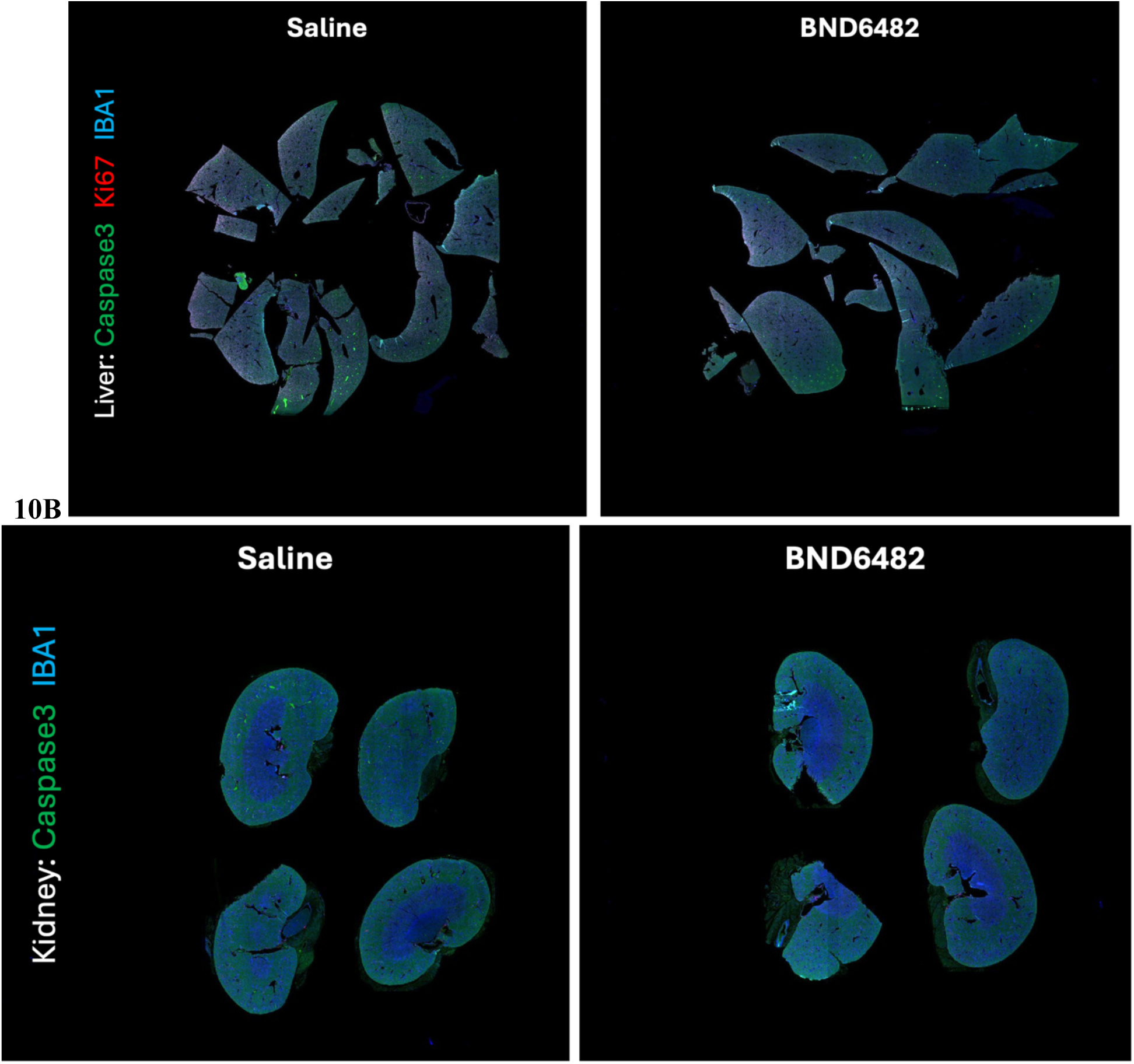
Liver and kidney toxicity assessment of BND6482 in EMT6 orthotopic mouse model. **(A)** Serum markers of liver and kidney toxicity after three IP injections of 5 mg/kg of BND6482 over 2 weeks. Error bars show s.e.m. **(B)** Examples of immunohistochemistry stain of liver and kidney injury markers, caspase 3, Ki67, and IBA1, after three IP injections of 5 mg/kg of BND6482.

## Conclusion and Discussion

Lipofection of the anti-miR-21 BNA-DNA-BNA gapmer BND5412 **(Fig. 2)** resulted in sub-nanomolar IC50 values for inhibiting proliferation across seven TNBC cell lines (**Fig. 5B**). IC50 values positively correlated with miR-21 copies/cell (**Fig. 5B**). Among the eight TNBC cell lines tested, MDA-MB-453 exhibited the weakest response to miR-21 inhibition by BND5412. This resistance is likely due to the fact that MDA-MB-453 is not a true triple negative cell line, as it harbors amplified HER2 signaling [91] and high androgen receptor (AR) expression [92]. These factors drive constitutive activation of HER2 and AR pathways, promoting sustained cell growth and survival. While miR-21 inhibition typically suppresses PI3K-AKT pathway by upregulating PDCD4, a downstream target of mTORC1, the persistent activation of HER2 and AR signaling enables continued activation of alternative AKT-driven survival pathways, ultimately diminishing the therapeutic efficacy of miR-21 inhibition. Overall, the miR-21 inhibitor BND5412 significantly slowed proliferation **(Fig. 5A)**, elevated tumor suppressor proteins **(Fig. 3C)**, reduced immune checkpoint proteins **(Fig. 4A-B)**, suppressed immune checkpoint gene expression **(Fig. 4C)**, and increased apoptosis **(Fig. 5C)** in multiple cell lines representing six genomic TNBC subtypes. These findings suggest that miR-21 blockade not only impairs proliferation and survival but also potentially enhances anti-tumor immunity in TNBC cells.

Comparing cumulative distributions of transcripts between miR-21 targets and all other genes showed a significant difference (**Fig. 6B**), indicating that BND5412 treatment of TNBC cells had global on-target effects on miR-21 regulated transcripts. No off-target change more than 4-fold was observed in any gene. RNA-seq enrichment analysis showed that miR-21 blockade activated interferon alpha and interferon gamma response, both of which have a positive effect on anti-tumor immunity [84, 85]. In addition, the *KRAS* inhibitory pathway was also significantly enriched in miR-21 BNA treated samples (**Fig. 6D**). Transcriptome analysis thus suggested additional unexpected anti-cancer benefits of miR-21 blockade.

Targeted delivery is the biggest challenge for RNA analog therapy. Current delivery approaches using liposomes or nanoparticles are suboptimal for long-term clinical use due to low delivery efficiency, toxicity, and restricted biodistribution to liver and kidneys [93]. Our designs provide safe, efficient, extra-hepatic delivery of nucleic acid therapeutics. Conjugating RNA analogs to receptor targeting peptide analogs (**Fig. 7B**) provides cell-type selective delivery. IGF1R is elevated in aggressive breast cancers, including TNBC [56].

We previously designed polyamide nucleic acid (PNA) (**Fig. 1**) oligomers with a protease-resistant retro-inverso cyclized D(CSKC) tetrapeptide IGF1 analog at the C-terminus of PNA to direct endocytosis into IGF1R-overexpressing cells [94, 95]. We found that radiolabeled PNA 12mers with a C-terminal cyclo-D(CSKC) stay in circulation by complexing with IGF1BP [96], extending the lifetime of the agent in the body, which reduces the necessary dose. Urine from mice injected with a [^99m^Tc]*MYCC* PNA-cyclo-D(CSKC) showed 83% of the radioactivity in an intact probe peak [97], illustrating *in vivo* stability.

We previously observed receptor-mediated PNA-cyclo-D(CSKC) knockdown of cyclin D1 protein in MCF7 breast cancer xenografts [98]. We have also used cyclo-D(CSKC) for tissue-specific delivery of various radioimaging agents. Thus, we were able to image *CCND1, MYCC, HER2,* or *KRAS2* mRNAs with nuclide-PNA-cyclo-D(CSKC) PET agents in mice bearing ER+ breast cancer xenografts [95, 98, 99], Her2+ breast cancer xenografts, illustrating DOX response by reduction of *HER2* mRNA PET SUV [100], pancreas cancer xenografts [99, 101], and transgenic mice with spontaneous mammary or lung tumors, illustrating response to cisplatin [102], with single mismatch specificity. Importantly, specific PET imaging was blocked by excess IGF1 [95].

The miR-21 BNA-DNA-BNA-peptide BND6482 presents a novel method to inhibit miR-21 by leveraging a peptide ligand for targeted delivery into TNBC cells without the need for lipofection. BND6482 demonstrated dose-dependent efficacy in high IGF1R-expressing HCC1806 cells, while showing no significant effect in low IGF1R-expressing MDA-MB-157 cells. In immunocompetent female Balb/c mice, intraperitoneal administration of AlexaFluor647-labeled BND6482 effectively localized to EMT6 allografts, leading to a reduction in tumor mass after 3 days of therapy. Further, a 2-week course of 5 mg/kg BND6482 halted tumor progression, resulting in a 74% mean reduction in tumor mass, with no observed liver or kidney toxicity, as confirmed by serum markers and histological analysis. Two-week treatment also reduced miR-21 levels and significantly upregulated PDCD4 mRNA, a miR-21 target, supporting the hypothesized mechanism of action. These findings indicate that BND6482 is an effective and safe therapeutic candidate for TNBC at the tested dose.

Using peptides as drug delivery molecules provides precise targeting, biocompatibility, and versatility, which are especially valuable in therapies where selective delivery to specific tissues or cells is critical, such as in cancer treatment [103]. Cell-penetrating peptides (CPPs), composed of basic amino acids or derived from protein translocation motifs, have been successfully conjugated to neutral backbone oligonucleotides such as phosphorodiamidate morpholino oligonucleotides (PMOs) and peptide nucleic acids (PNAs) for enhanced delivery [104]. However, conjugating CPPs to charged oligonucleotides has been less widely adopted due to the charge-charge interactions between the cationic CPPs and anionic oligonucleotides, which complicate synthesis and purification. To overcome these challenges, conjugating receptor-targeting peptides to charged oligonucleotides offers a promising alternative, as these peptide ligands facilitate cellular uptake via receptor-mediated endocytosis, bypassing the need for multiple cationic residues that are typically required for lipid bilayer destabilization. Recently, one study demonstrated a novel approach to targeted delivery of antisense oligonucleotides (ASOs) to pancreatic β-cells by conjugating ASOs to a GLP1R peptide agonist. The results show that this strategy enhances ASO uptake in pancreatic islets, leading to effective gene silencing in GLP1R-expressing cells both *in vitro* and *in vivo*, without significantly affecting gene expression in the liver or other non-target tissues [105]. Another study explored a novel approach for tumor-targeted gene silencing using systemic delivery of cRGD-conjugated siRNA, specifically targeting the VEGFR2 gene. The cRGD-siRNA conjugates were shown to selectively enter tumor cells expressing αvβ3 integrin, resulting in efficient gene knockdown, inhibition of angiogenesis, and significant reduction in tumor growth in both zebrafish and mouse models, without inducing immune responses or toxicity [106].

In this study, we present an RNA-peptide analog strategy targeting IGF1R for cancer cell-specific delivery, aimed at inhibiting proliferation and inducing apoptosis. Tumor suppression observed in a syngeneic mouse model highlights the potential of the anti-miR-21 RNA-peptide analog BND6482, supporting further preclinical proof-of-concept studies to facilitate its clinical translation.

## Materials and Methods

### BNA-DNA-BNA gapmers

Antisense BNA-DNA-BNA 5-5-5 gapmer phosphorothioates and peptide conjugates (**Table 1**) (Bio-Synthesis, Inc., Lewisville, TX 75057) were assembled by solid phase coupling of nucleotide phosphoramidites [4], 3’ addition of an alkyne linker, then click chemistry coupling of azidolysyl-peptides [107]. The gapmers and conjugates were analyzed by reversed phase liquid chromatography and MALDI-TOF mass spectroscopy as described [94, 108]. Calculated gapmer and conjugate masses agreed with manufacturer’s reported masses for BND5412: 5413.6 Da, BND5371: 5372.1 Da, BND6482: 6483.5 Da, and BND6372: 6371.6 Da.

### Cell lines and cell culture

Seven human TNBC cell lines, of European and African ancestry, which represent 80% of all TNBC subtypes [109], were studied: basal-like 1 MDA-MB-468 cells [110], basal-like 1 HCC1937 cells [111], basal-like 2 HCC1806 cells [111], mesenchymal stem-like MDA-MB-231 cells [112], mesenchymal stem–like MDA-MB-157 cells [109], mesenchymal stem–like MDA-MB-436 cells [109], immunomodulatory HCC1187 cells [111], mesenchymal BT549 cells [109], luminal androgen receptor MDA-MB-453 [109], and unclassified BT-20 cells [109], vs. a benign fibrocystic breast line, MCF10A [113], as a negative control. Two murine TNBC cell lines, EMT6 cells [114] and 4T1 cells [115], originating in female immunocompetent Balb/c mice, were studied. All cell lines were obtained from ATCC and maintained according to ATCC instructions.

### Cell treatment with BNA-DNA-BNA gapmers for western blot and real-time quantitative RT-PCR

TNBC cells were seeded in T-25 flask in complete medium without antibiotics the day before transfection. BNA-DNA-BNA gapmers were transfected into TNBC cells with Lipofectamine 3000 (Invitrogen) at a final gapmer concentration of 50 nM for 24-72 h at 37°C under 5% CO_2_, according to the manufacturer’s protocol. At the end of transfection, cells were detached and harvested with cell dissociation buffer.

### Dual-Glo luciferase reporter assay to determine IC50 for miR-21 inhibition

MDA-MB-231 and HCC1806 cells were seeded in 24-well plates to reach 50% confluency in complete medium without antibiotics the day before transfection. Cells were co-transfected with 100 ng of dual-luciferase psiCHECK2 plasmid (C8021, Promega) containing a perfect complement to miR-21 in the 3’UTR of the *Renilla* luciferase gene, and with a concentration gradient of BNA-DNA-BNA gapmers, using Lipofectamine 3000 (Invitrogen, CA). After 24 h, cells were lysed and Dual-Glo reporter assays were performed according to the manufacturer’s protocol (E2920, Promega). Luminescence intensity for each sample was measured with a Veritas Microplate Luminometer (Turner BioSystems), and each value from psiCHECK2 Renilla luciferase was normalized by psiCHECK2 firefly luciferase. IC50 was determined by the concentration at 50% elevation of the luciferase signal using non-linear regression in GraphPad Prism.

### Cell proliferation by Cell-Titer Glo assay after treatment with BNA-DNA-BNA gapmers

TNBC cells were seeded in 24-well plates to reach 50% confluency in complete medium without antibiotics the day before transfection. BNA-DNA-BNA gapmers were transfected into TNBC cells with Lipofectamine 3000 (Invitrogen) at a final gapmer concentration of 50 nM for 4 days for cell viability measurement over time, according to the manufacturer’s protocol. At the end of transfection, cellular viability was measured by quantitating lysed ATP using the Cell-Titer Glo Luminescent Cell Viability Assay (Promega). For cell proliferation IC50 measurements, TNBC cells seeded in 24-well plates were transfected with concentration gradients of BNA-DNA-BNA gapmers for 48 h, followed by CellTiter-Glo Luminescent Cell Viability Assay (Promega).

Luminescence intensity for each sample was measured with a BioTek Synergy HT Reader.

### Western blots of cellular proteins in treated cells

Treated cells were detached by cell dissociation buffer and harvested with 1×PBS, then lysed in cell lysis buffer (Invitrogen) with protease inhibitor cocktail (P-2714, Sigma). Lysate protein concentrations were quantified by the Bradford Assay (Bio-Rad). Lysate aliquots containing 30 µg protein were separated on NuPAGE 4-12% Bis-Tris gels (Invitrogen), transferred to PVDF membranes, blocked with blocking buffer, and incubated with antibodies against PDCD4 protein (ab80590, Abcam), PTEN protein (9552S, Cell Signaling), CD47 protein (AF4670, R&D Systems), PD-L1 protein (501124188, ThermoFisher), PD-L2 protein (501123155, ThermoFisher), Jak2 protein (D2E12, Cell Signaling), and β-actin (AM4302, Ambion), followed by incubation with secondary antibodies labeled with horseradish peroxidase (Invitrogen). The resulting protein bands were imaged by luminescence using a SuperSignal West Femto Chemiluminescent Substrate (Thermo Scientific), on a Bio-Rad ChemiDoc Imaging System, and analyzed with Image Lab software (Bio-Rad).

### Lactate dehydrogenase (LDH) assay to detect apoptosis in treated cells

Cells in 24-well plates were treated with 50 nM BNA-DNA-BNA gapmers as above. 72 h later, induction of apoptosis was deduced from LDH released into the medium due to cell lysis, using CytoTox 96® Non-Radioactive Cytotoxicity Assay (Promega).

### Real-Time PCR

Total RNA from treated TNBC cells was extracted using a mirVana miRNA isolation kit (ThermoFishser) according to the manufacturer’s protocol. Total RNA from treated EMT6 tumors was extracted using TRIzol Plus RNA Purification Kit (ThermoFishser) according to the manufacturer’s protocol. For qPCR of miRNAs, 100 ng of purified total RNA were reverse transcribed with a TaqMan miRNA reverse transcription kit (#4366597, ThermoFisher). qPCR of miRNAs was performed with a miRNA Gene Expression Assay (ThermoFisher) on a Applied Biosystems StepOnePlus RT-PCT system. Primers specific for miR-21 (Assay ID 000397) and internal control RNA U6 (Assay ID 001973) for both reverse transcription and qPCR were obtained from ThermoFisher. The average absolute values of triplicate samples for the same miRNA were calculated and normalized to U6 RNA, measured by the comparative Ct (2^-ΔΔCt^) method [116].

For qPCR of tumor suppressor and immune checkpoint mRNAs, 500 ng of purified total RNA were reverse transcribed with a High Capacity cDNA Reverse Transcription Kit (ThermoFisher, cat#4368814). qPCR of mRNAs was performed using TaqMan Gene Expression Assay (ThermoFisher) on a Applied Biosystems StepOnePlus RT-PCT system with the following TaqMan primers/probes to detect transcripts: *GAPDH* (Hs02786624_g1); *PDCD4* (Hs00377253_m1); *CD47* (Hs00179953_m1); *PDL1* (Hs00204257_m1); *PDL2* (Hs00228839_m1); *JAK2* (Hs01078136_m1). The average of triplicate samples for the same mRNA was calculated and normalized to the internal control gene *GAPDH* (Hs03929097_g1), by the comparative Ct (2^-ΔΔCt^) method [116].

To quantify the copy number of miR-21 per cell, synthetic miR-21 RNA was reverse transcribed into cDNA using the TaqMan miRNA Reverse Transcription Kit (#4366597, ThermoFisher). A standard curve was generated by performing qPCR on five concentrations of synthetic miR-21, ranging from 1.8×10^7 to 3×10×8 copies/L, using the miR-21 TaqMan Gene Expression Assay (Assay ID 000397, ThermoFisher). In parallel, total RNA extracted from cells underwent miR-21 qPCR using the same assay. The total miR-21 copy number in each RNA sample was then calculated based on the standard curve. Finally, the copy number per cell was determined by dividing the total miR-21 copy number by the number of cells used for RNA extraction.

### RNA-seq measurements of transcriptome changes of miR-21 target mRNAs in treated cells: Total

RNA was extracted from HCC1806 cells 24 hours post-transfection with EC90 (10.7 nM) of BND5412, using the mirVana miRNA isolation kit (ThermoFisher) following the manufacturer’s protocol. RNA quality assessment, library preparation using the Stranded Total RNA Prep, Ligation with Ribo-Zero Plus kit (Illumina), and RNA sequencing (RNA-seq) via NovaSeq 6000 (Illumina), generating 60 million reads per sample in a 2×100 bp paired-end configuration, were conducted at the Cancer Genomics & Bioinformatics Core at Thomas Jefferson University. Samples were multiplexed across two flow cell lanes. FASTQ files were aligned using STAR, with quality checks performed via RSeQC, and aligned reads counted with HTSeq.

Differential gene expression analysis was conducted with DESeq2, and Jensen Disease enrichment analysis was performed using the iDEP platform 2.01 (http://bioinformatics.sdstate.edu/idep96/). Gene Set Enrichment Analysis (GSEA) was performed using GSEA software version 4.3.2 from the Broad Institute to evaluate the enrichment of predefined gene sets in differentially expressed genes from RNA-seq data. Differentially expressed genes were identified by comparing control and treated samples using DESeq2, with adjusted *p*-value < 0.05 as thresholds for significance. The ranked gene list was generated based on the log2 fold change of gene expression between treated and control samples. This ranked list was used as input for GSEA. The analysis was conducted using the Hallmark gene sets from the Molecular Signatures Database (MSigDB).

### Confocal fluorescent microscopy

HCC1806 cells were seeded at a density of 5 × 10^4 cells/ml in 2 ml of growth medium in Ibidi 35 mm dishes and cultured overnight at 37°C in a humidified atmosphere containing 5% CO_2_. Following the initial incubation, cells were treated with 100 nM Cy5-BND6482 in fresh growth medium for 4 hours under identical conditions. Subsequently, the cells were incubated with Lysotracker Green (80 nM final concentration) for 30 minutes at 37°C. After incubation, the cells were gently washed thrice with Ca2+ and Mg2+ containing phosphate-buffered saline (PBS) and immersed in 2 ml of imaging medium. Live-cell confocal microscopy was performed using a Nikon A1R HD confocal microscope equipped with a 60× objective lens and 3× optical zoom

### Serum stability of BND6482

BND6482 was incubated in either 50% or 90% FBS, diluted with RPMI 1640 medium at 37°C, to a final concentration of 2 µM. Aliquots (10 µL) were collected at specified time points (0, 0.5, 1, 2, 4, 8, 24, and 48 hours) and stored at −80°C until analysis. After the incubation period, all samples were resolved using a 15% TBE-Urea gel with TBE running buffer. The gel was run at a constant voltage of 180V for 50 minutes, followed by staining in 50 mL of 1x SYBR Gold Nucleic Acid Gel Stain (ThermoFisher, S11494), diluted in TBE running buffer, for 25 minutes. The stained gel was imaged using an iBright FL1500 Imaging System (Invitrogen).

### Orthotopic TNBC allograft treatment

Murine EMT6 TNBC cells [114], originating in female immunocompetent Balb/c mice, express high miR-21 and median IGF1R relative to other TNBC cells [114]. Therefore, we chose EMT6 cells to create an orthotopic female immunocompetent Balb/c model for TNBC tumor progression. In groups of 5 syngeneic, immunocompetent 4-8 week old female Balb/c mice, we implanted 1×10^5^ EMT6 cells into a mammary fat pad of each subject under isoflurane anesthesia. Tumors were allowed to grow to 5 mm diameter. Then we administered BND6482, scrambled control BND6372, Trodelvy, PBS, or saline vehicle intraperitoneally to each subject, at 5 mg/kg twice a week on days 0, 3, 7, and 10 over 13 days. Palpable tumor volume were measured using calipers with the formular (0.5 x length x width x width), along with dissected tumor weights.

### Fluorescent imaging of biodistribution and tumor uptake

To assess biodistribution, EMT6 tumor-bearing mice received a single intraperitoneal (IP) injection of 5 mg/kg AlexaFluor647-labeled BND6482 at time 0.

Mice were then positioned prone under isoflurane anesthesia and imaged using an In-Vivo Multispectral FX PRO optical imaging system. Fluorescence and X-ray images were captured at 2, 24, 48, and 96 hours post-injection.

For tumor uptake analysis, EMT6 tumors were allowed to grow to approximately 5 mm in diameter in 4-to 8-week-old female Balb/c mice. Mice were then administered either a saline vehicle control or AlexaFluor647-labeled BND6482 at 5 mg/kg via IP injection once daily for three consecutive days. On the fourth day, tumors were harvested and imaged using the AlexaFluor647 fluorescence channel on a Bio-Rad ChemiDoc Imaging System.

### Serum markers of liver and kidney toxicity

Serum markers of liver and kidney toxicity were assessed from blood collected immediately following sacrifice. The blood was drawn after three IP injections of 5 mg/kg BND6482, administered over a 2-week period on days 0, 6, and 12, once the tumors had reached a diameter of 5 mm.

### Histological analysis of thin sections of liver, and kidney

Histological analysis of the liver and kidneys was conducted following three IP injections of 5 mg/kg BND6482, administered over a 2-week period on days 0, 6, and 12, once tumors had reached a diameter of 5 mm. Liver and kidney tissues were fixed in 10% neutral buffered formalin, embedded in paraffin, and processed for immunofluorescence staining by the Comparative Pathology Core at the University of Pennsylvania School of Veterinary Medicine.

### Statistical Analysis

All experimental measurements were performed independently at least three times. Significance was assessed by Student’s t-test. One-way ANOVA and Kolmogorov-Smirnov tests were conducted with GraphPad Prism 10. Prior to applying one-way ANOVA and t-tests, data normality was assessed using the Shapiro-Wilk test.

## Acknowledgments

We thank the late Dr. Edith Mitchell for advice on clinical translation.

This work was supported by NIH STTR grant 1 R01 CA235707 to E.W., and by Bound Therapeutics LLC

## Abbreviations

BNA: bridged nucleic acid
CD47: cluster of differentiation 47
IGF1: insulin-like growth factor 1
Jak2: Janus kinase 2
LNA: locked nucleic acid
BNA: bridged nucleic acid
miRNA: micro ribonucleic acid
miR-21: microRNA-21
oncomiR: oncogenic miRNA
PBS: phosphate buffered saline
PDCD4: programmed cell death 4
PD-L1: programmed death-ligand 1
PD-L2: programmed death-ligand 2
PTEN: phosphatase and tensin homolog
RNA: ribonucleic acid
TNBC: triple negative breast cancer
ER: estrogen receptor
PR: progesterone receptor
PARP: poly(ADP-ribose) polymerase
RTK: receptor tyrosine kinase
ALP: alkaline phosphatase
ALT: alanine aminotransferase
AST: aspartate aminotransferase

## References

1. Koziolkiewicz, M., et al., Stereodifferentiation--the effect of P chirality of oligo(nucleoside phosphorothioates) on the activity of bacterial RNase H. Nucleic Acids Res, 1995. 23(24): p. 5000–5.

2. Miller, P.S., et al., Control of ribonucleic acid function by oligonucleoside methylphosphonates. Biochimie, 1985. 67(7-8): p. 769–76.

3. Vester, B. and J. Wengel, LNA (locked nucleic acid): high-affinity targeting of complementary RNA and DNA. Biochemistry, 2004. 43(42): p. 13233–41.

4. Rahman, S.M., et al., Design, synthesis, and properties of 2’,4’-BNA(NC): a bridged nucleic acid analogue. J Am Chem Soc, 2008. 130(14): p. 4886–96.

5. American_Cancer_Society. *Breast Cancer Facts and Figures*. 2020; Available from: https://www.cancer.org/research/cancer-facts-statistics/breast-cancer-facts-figures.html.

6. Nielsen, P.E., Gene targeting and expression modulation by peptide nucleic acids (PNA). Curr Pharm Des, 2010. 16(28): p. 3118–23.

7. Kalimutho, M., et al., Targeted therapies for triple-negative breast cancer: combating a stubborn disease. Trends Pharmacol Sci, 2015. 36(12): p. 822–46.

8. Hudis, C.A. and L. Gianni, Triple-negative breast cancer: an unmet medical need. Oncologist, 2011. 16 **Suppl 1**: p. 1–11.

9. Mitchell, E.P., Triple-negative breast cancer: How can outcomes be improved? Frontline, 2015(Fall): p. 5.

10. Dietze, E.C., et al., Triple-negative breast cancer in African-American women: disparities versus biology. Nat Rev Cancer, 2015. 15(4): p. 248–54.

11. Miller, T.W., et al., A gene expression signature from human breast cancer cells with acquired hormone independence identifies MYC as a mediator of antiestrogen resistance. Clinical Cancer Research, 2011. 17(7): p. 2024–2034.

12. Musgrove, E.A. and R.L. Sutherland, Biological determinants of endocrine resistance in breast cancer. Nat Rev Cancer, 2009. 9(9): p. 631–43.

13. Syed, B.A., The breast cancer market. Nat Rev Drug Discov, 2015. 14(4): p. 233–4.

14. Perez, E.A., et al., Etirinotecan pegol (NKTR-102) versus treatment of physician’s choice in women with advanced breast cancer previously treated with an anthracycline, a taxane, and capecitabine (BEACON): a randomised, open-label, multicentre, phase 3 trial. Lancet Oncol, 2015. 16(15): p. 1556–68.

15. Merck_Sharp_&_Dohme. Study of pembrolizumab (MK-3475) monotherapy for metastatic triple-negative breast cancer (MK-3475-086/KEYNOTE-086). NCT02447003 2015; Available from: https://clinicaltrials.gov/ct2/show/NCT02447003?term=pembrolizumab&cond=%22Triple+Negative+Breast+Neoplasms%22&rank=4.

16. Yardley, D.A., et al., EMERGE: A Randomized Phase II Study of the Antibody-Drug Conjugate Glembatumumab Vedotin in Advanced Glycoprotein NMB-Expressing Breast Cancer. J Clin Oncol, 2015. 33(14): p. 1609–19.

17. Luo, L. and K. Keyomarsi, PARP inhibitors as single agents and in combination therapy: the most promising treatment strategies in clinical trials for BRCA-mutant ovarian and triple-negative breast cancers. Expert Opin Investig Drugs, 2022. 31(6): p. 607–631.

18. Bagegni, N.A., et al., Targeted Treatment for High-Risk Early-Stage Triple-Negative Breast Cancer: Spotlight on Pembrolizumab. Breast Cancer (Dove Med Press), 2022. 14: p. 113–123.

19. Amin, D.N., et al., Resiliency and vulnerability in the HER2-HER3 tumorigenic driver. Sci Transl Med, 2010. 2(16): p. 16ra7.

20. Virgil, H. Atezolizumab TNBC Indication Withdrawn By Manufacturer After Talks With FDA. 2021; Available from: https://www.cancernetwork.com/view/atezolizumab-tnbc-indication-withdrawn-by-manufacturer-after-talks-with-fda.

21. Rugo, H.S., et al., Overall survival with sacituzumab govitecan in hormone receptor-positive and human epidermal growth factor receptor 2-negative metastatic breast cancer (TROPiCS-02): a randomised, open-label, multicentre, phase 3 trial. Lancet, 2023. 402(10411): p. 1423–1433.

22. Bartel, D.P., MicroRNAs: target recognition and regulatory functions. Cell, 2009. 136(2): p. 215–33.

23. Calin, G.A. and C.M. Croce, MicroRNA signatures in human cancers. Nature reviews. Cancer, 2006. 6(11): p. 857–66.

24. Croce, C.M., Causes and consequences of microRNA dysregulation in cancer. Nat Rev Genet, 2009. 10(10): p. 704–14.

25. Di Leva, G., M. Garofalo, and C.M. Croce, MicroRNAs in cancer. Annu Rev Pathol, 2014. 9: p. 287–314.

26. Sanders, J.M., et al., Effects of hypoxanthine substitution in peptide nucleic acids targeting KRAS2 oncogenic mRNA molecules: theory and experiment. Journal of Physical Chemistry B, 2013. 117(39): p. 11584–11595.

27. Iorio, M.V., et al., MicroRNA gene expression deregulation in human breast cancer. Cancer Res, 2005. 65(16): p. 7065–70.

28. Radojicic, J., et al., MicroRNA expression analysis in triple-negative (ER, PR and Her2/neu) breast cancer. Cell Cycle, 2011. 10(3): p. 507–17.

29. MacKenzie, T.A., et al., Stromal expression of miR-21 identifies high-risk group in triple-negative breast cancer. Am J Pathol, 2014. 184(12): p. 3217–25.

30. Dong, G., et al., High expression of miR-21 in triple-negative breast cancers was correlated with a poor prognosis and promoted tumor cell in vitro proliferation. Medical Oncology, 2014. 31(7): p. 1–10.

31. Zhang, Z.J. and S.L. Ma, miRNAs in breast cancer tumorigenesis (Review). Oncol Rep, 2012. 27(4): p. 903–10.

32. Corcoran, C., et al., Intracellular and extracellular microRNAs in breast cancer. Clin Chem, 2011. 57(1): p. 18–32.

33. Ozgun, A., et al., MicroRNA-21 as an indicator of aggressive phenotype in breast cancer. Onkologie, 2013. 36(3): p. 115–8.

34. Anastasov, N., et al., Radiation resistance due to high expression of miR-21 and G2/M checkpoint arrest in breast cancer cells. Radiat Oncol, 2012. 7: p. 206.

35. Min, W., et al., The expression and significance of five types of miRNAs in breast cancer. Med Sci Monit Basic Res, 2014. 20: p. 97–104.

36. Najjary, S., et al., Role of miR-21 as an authentic oncogene in mediating drug resistance in breast cancer. Gene, 2020. 738: p. 144453.

37. Shao, B., et al., Plasma microRNAs Predict Chemoresistance in Patients With Metastatic Breast Cancer. Technol Cancer Res Treat, 2019. 18: p. 1533033819828709.

38. Dan, T., et al., miR-21 Plays a Dual Role in Tumor Formation and Cytotoxic Response in Breast Tumors. Cancers (Basel), 2021. 13(4).

39. Lu, M., et al., An analysis of human microRNA and disease associations. PLoS ONE, 2008. 3(10): p. e3420.

40. Meng, F., et al., MicroRNA-21 regulates expression of the PTEN tumor suppressor gene in human hepatocellular cancer. Gastroenterology, 2007. 133(2): p. 647–58.

41. Sayed, D., et al., MicroRNA-21 targets Sprouty2 and promotes cellular outgrowths. Mol Biol Cell, 2008. 19(8): p. 3272–82.

42. Zhu, S., et al., MicroRNA-21 targets tumor suppressor genes in invasion and metastasis. Cell Res, 2008. 18(3): p. 350–9.

43. Zhu, S., et al., MicroRNA-21 targets the tumor suppressor gene tropomyosin 1 (TPM1). J Biol Chem, 2007. 282(19): p. 14328–36.

44. Chen, Y., et al., Loss of PDCD4 expression in human lung cancer correlates with tumour progression and prognosis. J Pathol, 2003. 200(5): p. 640–6.

45. Lee, S., et al., Differential expression in normal-adenoma-carcinoma sequence suggests complex molecular carcinogenesis in colon. Oncol Rep, 2006. 16(4): p. 747–54.

46. Ma, G., et al., [Expression of programmed cell death 4 and its clinicopathological significance in human pancreatic cancer]. Zhongguo Yi Xue Ke Xue Yuan Xue Bao, 2005. 27(5): p. 597–600.

47. Jansen, A.P., et al., Characterization of programmed cell death 4 in multiple human cancers reveals a novel enhancer of drug sensitivity. Mol Cancer Ther, 2004. 3(2): p. 103–10.

48. Suzuki, C., et al., PDCD4 inhibits translation initiation by binding to eIF4A using both its MA3 domains. Proc Natl Acad Sci U S A, 2008. 105(9): p. 3274–9.

49. Lankat-Buttgereit, B. and R. Goke, The tumour suppressor Pdcd4: recent advances in the elucidation of function and regulation. Biol Cell, 2009. 101(6): p. 309–17.

50. Maehama, T., PTEN: its deregulation and tumorigenesis. Biol Pharm Bull, 2007. 30(9): p. 1624–7.

51. Bailey, C.M., et al., Biological functions of maspin. J Cell Physiol, 2006. 209(3): p. 617–24.

52. Loffler, D., et al., Interleukin-6 dependent survival of multiple myeloma cells involves the Stat3-mediated induction of microRNA-21 through a highly conserved enhancer. Blood, 2007. 110(4): p. 1330–3.

53. Yang, C.H., et al., IFN induces miR-21 through a signal transducer and activator of transcription 3-dependent pathway as a suppressive negative feedback on IFN-induced apoptosis. Cancer Res, 2010. 70(20): p. 8108–16.

54. Bhat, B. and E. Marcusson, MicroRNA compounds and methods for modulating miR-21 activity, 8969317, Editor. 2015, Regulus Therapeutics: USA. p. 69.

55. Li, J., et al., Structural basis of the activation of type 1 insulin-like growth factor receptor. Nat Commun, 2019. 10(1): p. 4567.

56. Surmacz, E., Function of the IGF-I receptor in breast cancer. J Mammary Gland Biol Neoplasia, 2000. 5(1): p. 95–105.

57. Law, J.H., et al., Phosphorylated insulin-like growth factor-i/insulin receptor is present in all breast cancer subtypes and is related to poor survival. Cancer Res, 2008. 68(24): p. 10238–46.

58. Yerushalmi, R., et al., Insulin-like growth factor receptor (IGF-1R) in breast cancer subtypes. Breast Cancer Res Treat, 2012. 132(1): p. 131–42.

59. Tian, X., et al., External imaging of CCND1, MYC, and KRAS oncogene mRNAs with tumor-targeted radionuclide-PNA-peptide chimeras. Annals of the New York Academy of Sciences, 2005. 1059(1): p. 106–144.

60. Dobre, M., et al., Dysregulation of miRNAs Targeting the IGF-1R Pathway in Pancreatic Ductal Adenocarcinoma. Cells, 2021. 10(8).

61. Miyashita, K., et al., N-Methyl substituted 2’,4’-BNANC: a highly nuclease-resistant nucleic acid analogue with high-affinity RNA selective hybridization. Chem Commun (Camb), 2007(36): p. 3765–7.

62. Torigoe, H., et al., 2’-O,4’-C-aminomethylene-bridged nucleic acid modification with enhancement of nuclease resistance promotes pyrimidine motif triplex nucleic acid formation at physiological pH. Chemistry, 2011. 17(9): p. 2742–51.

63. Yamamoto, T., et al., Cholesterol-lowering Action of BNA-based Antisense Oligonucleotides Targeting PCSK9 in Atherogenic Diet-induced Hypercholesterolemic Mice. Mol Ther Nucleic Acids, 2012. 1: p. e22.

64. Mukai, H., et al., Quantitative evaluation of the improvement in the pharmacokinetics of a nucleic acid drug delivery system by dynamic PET imaging with 18F-incorporated oligodeoxynucleotides. Journal of Controlled Release, 2014. 180: p. 92–99.

65. Manning, K.S., et al., BNA(NC) Gapmers Revert Splicing and Reduce RNA Foci with Low Toxicity in Myotonic Dystrophy Cells. ACS Chem Biol, 2017. 12(10): p. 2503–2509.

66. Jepsen, J.S., M.D. Sorensen, and J. Wengel, Locked nucleic acid: a potent nucleic acid analog in therapeutics and biotechnology. Oligonucleotides, 2004. 14(2): p. 130–46.

67. Elmen, J., et al., LNA-mediated microRNA silencing in non-human primates. Nature, 2008. 452(7189): p. 896–9.

68. Elmen, J., et al., Antagonism of microRNA-122 in mice by systemically administered LNA-antimiR leads to up-regulation of a large set of predicted target mRNAs in the liver. Nucleic Acids Res, 2008. 36(4): p. 1153–62.

69. Obad, S., et al., Silencing of microRNA families by seed-targeting tiny LNAs. Nat Genet, 2011. 43(4): p. 371–8.

70. Swayze, E.E., et al., Antisense oligonucleotides containing locked nucleic acid improve potency but cause significant hepatotoxicity in animals. Nucleic Acids Res, 2007. 35(2): p. 687–700.

71. Kasuya, T., et al., Ribonuclease H1-dependent hepatotoxicity caused by locked nucleic acid-modified gapmer antisense oligonucleotides. Sci Rep, 2016. 6: p. 30377.

72. Kakiuchi-Kiyota, S., et al., Comparison of hepatic transcription profiles of locked ribonucleic acid antisense oligonucleotides: evidence of distinct pathways contributing to non-target mediated toxicity in mice. Toxicol Sci, 2014. 138(1): p. 234–48.

73. Helene, C. and J.J. Toulme, Specific regulation of gene expression by antisense, sense and antigene nucleic acids. Biochim Biophys Acta, 1990. 1049(2): p. 99–125.

74. Jin, Y.-Y., J. Andrade, and E. Wickstrom, Non-specific blocking of miR-17-5p guide strand in triple negative breast cancer cells by amplifying passenger strand activity. PLoS One, 2015. 10(12): p. e0142574.

75. Frankel, L.B., et al., Programmed cell death 4 (PDCD4) is an important functional target of the microRNA miR-21 in breast cancer cells. J Biol Chem, 2008. 283(2): p. 1026–33.

76. Paladini, L., et al., Targeting microRNAs as key modulators of tumor immune response. J Exp Clin Cancer Res, 2016. 35: p. 103.

77. Chi, L.H., et al., MicroRNA-21 is immunosuppressive and pro-metastatic via separate mechanisms. Oncogenesis, 2022. 11(1): p. 38.

78. Jiang, Z., et al., Targeting CD47 for cancer immunotherapy. J Hematol Oncol, 2021. 14(1): p. 180.

79. Deng, H., et al., New hope for tumor immunotherapy: the macrophage-related “do not eat me” signaling pathway. Front Pharmacol, 2023. 14: p. 1228962.

80. Hudson, K., et al., The Extrinsic and Intrinsic Roles of PD-L1 and Its Receptor PD-1: Implications for Immunotherapy Treatment. Front Immunol, 2020. 11: p. 568931.

81. Huang, B., X. Lang, and X. Li, The role of IL-6/JAK2/STAT3 signaling pathway in cancers. Front Oncol, 2022. 12: p. 1023177.

82. Owen, K.L., N.K. Brockwell, and B.S. Parker, JAK-STAT Signaling: A Double-Edged Sword of Immune Regulation and Cancer Progression. Cancers (Basel), 2019. 11(12).

83. Yin, L., et al., Triple-negative breast cancer molecular subtyping and treatment progress. Breast Cancer Res, 2020. 22(1): p. 61.

84. Jorgovanovic, D., et al., Roles of IFN-gamma in tumor progression and regression: a review. Biomark Res, 2020. 8: p. 49.

85. Vidal, P., Interferon alpha in cancer immunoediting: From elimination to escape. Scand J Immunol, 2020. 91(5): p. e12863.

86. Pietrzkowski, Z., et al., Inhibition of cellular proliferation by peptide analogues of insulin-like growth factor 1. Cancer Res, 1992. 52(23): p. 6447–6451.

87. Jin, Y.-Y. and E. Wickstrom, Specific blocking of miR-17-5p guide strand in triple negative breast cancer cells, without amplifying passenger strand activity. Cancer Research, 2016. 76(S3): p. B41.

88. Cesarone, G., et al., Insulin receptor substrate 1 knockdown in human MCF7 estrogen receptor-positive breast cancer cells by nuclease-resistant IRS1 siRNA conjugated to a disulfide-bridged D-peptide analog of insulin-like growth factor 1. Bioconjugate Chemistry, 2007. 18(6): p. 1831–1840.

89. de Lint, K., et al., Sensitizing Triple-Negative Breast Cancer to PI3K Inhibition by Cotargeting IGF1R. Mol Cancer Ther, 2016. 15(7): p. 1545–56.

90. Romero-Palomo, F., et al., *S*afety, Tissue Distribution, and Metabolism of LNA-Containing Antisense Oligonucleotides in Rats. Toxicol Pathol, 2021. 49(6): p. 1174–1192.

91. Rahmati, M., et al., Investigating the cytotoxic and anti-proliferative effects of trastuzumab on MDA-MB-453 and MDA-MB-468 breast cell lines with different levels of HER2 expression. Journal of Applied Biotechnology Reports, 2020. 7(2): p. 87–92.

92. Hall, R.E., et al., MDA-MB-453, an androgen-responsive human breast carcinoma cell line with high level androgen receptor expression. Eur J Cancer, 1994. 30A(4): p. 484–90.

93. Juliano, R.L., The delivery of therapeutic oligonucleotides. Nucleic Acids Res, 2016. 44(14): p. 6518–48.

94. Basu, S. and E. Wickstrom, Synthesis and characterization of a peptide nucleic acid conjugated to a D-peptide analog of insulin-like growth factor 1 for increased cellular uptake. Bioconjugate Chemistry, 1997. 8(4): p. 481–488.

95. Tian, X., et al., PET imaging of CCND1 mRNA in human MCF7 estrogen receptor-positive breast cancer xenografts with an oncogene-specific [^64^Cu]DO3A-PNA-peptide radiohybridization probe. Journal of Nuclear Medicine, 2007. 48(10): p. 1699–1707.

96. Opitz, A.W., et al., Physiologically based pharmacokinetics of molecular imaging nanoparticles for mRNA detection determined in tumor-bearing mice. Oligonucleotides, 2010. 20(3): p. 117–25.

97. Tian, X., et al., Imaging oncogene expression. Annals of the New York Academy of Sciences, 2003. 1002: p. 165–188.

98. Tian, X., et al., External imaging of CCND1 cancer gene activity in experimental human breast cancer xenografts with Tc-99m-peptide-PNA-peptide chimeras. Journal of Nuclear Medicine, 2004. 45(12): p. 2070–2082.

99. Tian, X., et al., External imaging of CCND1, MYC, and KRAS oncogene mRNAs with tumor-targeted radionuclide-PNA-peptide chimeras. Annals of the New York Academy of Sciences, 2005. 1059: p. 106–44.

100. Paudyal, B., et al., Determining efficacy of breast cancer therapy by PET imaging of HER2 mRNA. Nuclear Medicine and Biology, 2013. 40(8): p. 994–999.

101. Chakrabarti, A., et al., Radiohybridization PET imaging of KRAS G12D mRNA expression in human pancreas cancer xenografts with [^64^Cu]DO3A-peptide nucleic acid-peptide nanoparticles. Cancer Biology & Therapy, 2007. 6(6): p. 948–956.

102. Chen, C.-P., et al., Hypoxanthine-containing PNA probes for imaging the mutations of KRAS2 oncogene mRNA, in XX International Roundtable on Nucleosides, Nucleotides, and Nucleic Acids. 2012: Montreal, Canada. p. 150.

103. Cooper, B.M., et al., Peptides as a platform for targeted therapeutics for cancer: peptide-drug conjugates (PDCs). Chem Soc Rev, 2021. 50(3): p. 1480–1494.

104. Yin, H., et al., Cell-penetrating peptide-conjugated antisense oligonucleotides restore systemic muscle and cardiac dystrophin expression and function. Hum Mol Genet, 2008. 17(24): p. 3909–18.

105. Ammala, C., et al., Targeted delivery of antisense oligonucleotides to pancreatic beta-cells. Sci Adv, 2018. 4(10): p. eaat3386.

106. Liu, X., et al., Tumor-targeted in vivo gene silencing via systemic delivery of cRGD-conjugated siRNA. Nucleic Acids Res, 2014. 42(18): p. 11805–17.

107. Gogoi, K., et al., A versatile method for the preparation of conjugates of peptides with DNA/PNA/analog by employing chemo-selective click reaction in water. Nucleic Acids Res, 2007. 35(21): p. e139.

108. Tian, X. and E. Wickstrom, Continuous solid-phase synthesis and disulfide cyclization of peptide-PNA-peptide chimeras. Organic Letters, 2002. 4(23): p. 4013–4016.

109. Lehmann, B.D., et al., Identification of human triple-negative breast cancer subtypes and preclinical models for selection of targeted therapies. J Clin Invest, 2011. 121(7): p. 2750–67.

110. Cailleau, R., M. Olive, and Q.V. Cruciger, Long-term human breast carcinoma cell lines of metastatic origin: preliminary characterization. In Vitro, 1978. 14(11): p. 911–5.

111. Gazdar, A.F., et al., Characterization of paired tumor and non-tumor cell lines established from patients with breast cancer. Int J Cancer, 1998. 78(6): p. 766–74.

112. Neve, R.M., et al., A collection of breast cancer cell lines for the study of functionally distinct cancer subtypes. Cancer Cell, 2006. 10(6): p. 515–27.

113. Soule, H.D., et al., Isolation and characterization of a spontaneously immortalized human breast epithelial cell line, MCF-10. Cancer Res, 1990. 50(18): p. 6075–86.

114. Trettel, F., et al., Dominant phenotypes produced by the HD mutation in STHdh(Q111) striatal cells. Hum Mol Genet, 2000. 9(19): p. 2799–809.

115. Pulaski, B.A. and S. Ostrand-Rosenberg, Reduction of established spontaneous mammary carcinoma metastases following immunotherapy with major histocompatibility complex class II and B7.1 cell-based tumor vaccines. Cancer Res, 1998. 58(7): p. 1486–93.

116. Livak, K.J. and T.D. Schmittgen, Analysis of relative gene expression data using real-time quantitative PCR and the 2(-Delta Delta C(T)) Method. Methods, 2001. 25(4): p. 402–8.

